# Neonatal genetics of gene expression reveal the origins of autoimmune and allergic disease risk

**DOI:** 10.1101/683086

**Authors:** Qin Qin Huang, Howard H. F. Tang, Shu Mei Teo, Scott C. Ritchie, Artika P. Nath, Marta Brozynska, Agus Salim, Andrew Bakshi, Barbara J. Holt, Danny Mok, Chiea Chuen Khor, Peter D. Sly, Patrick G. Holt, Kathryn E. Holt, Michael Inouye

**Affiliations:** Cambridge Baker Systems Genomics Initiative, Baker Heart and Diabetes Institute, Melbourne, Victoria 3004, Australia; Department of Clinical Pathology, University of Melbourne, Parkville, Victoria 3010, Australia; School of BioSciences, The University of Melbourne, Parkville, Victoria 3010, Australia; Cambridge Baker Systems Genomics Initiative, Department of Public Health and Primary Care, University of Cambridge, Cambridge CB1 8RN, United Kingdom; Baker Heart and Diabetes Institute, Melbourne, Victoria 3004, Australia; Department of Mathematics and Statistics, School of Engineering and Mathematical Sciences, La Trobe University, Bundoora, Victoria 3086, Australia; Monash Biomedicine Discovery Institute, Prostate Cancer Research Group, Department of Anatomy and Developmental Biology, Monash University, Clayton, Victoria 3800, Australia; Telethon Kids Institute, The University of Western Australia, Perth, Western Australia 6009, Australia; Human Genetics, Genome Institute of Singapore, Agency for Science, Technology and Research, Singapore 138672, Singapore; Child Health Research Centre, The University of Queensland, Brisbane, Queensland 4101, Australia; Department of Infectious Diseases, Central Clinical School, Monash University, Melbourne, Victoria 3004, Australia; The London School of Hygiene and Tropical Medicine, London WC1E 7TH, United Kingdom; The Alan Turing Institute, London, United Kingdom

## Abstract

Chronic immune-mediated diseases of adulthood often originate in early childhood. To investigate genetic associations between neonatal immunity and disease, we collected cord blood samples from a birth cohort and mapped expression quantitative trait loci (eQTLs) in resting monocytes and CD4^+^ T cells as well as in response to lipopolysaccharide (LPS) or phytohemagglutinin (PHA) stimulation, respectively. *Cis*-eQTLs were largely specific to cell type or stimulation, and response eQTLs were identified for 31% of genes with *cis*-eQTLs (eGenes) in monocytes and 52% of eGenes in CD4^+^ T cells. We identified *trans*-eQTLs and mapped *cis* regulatory factors which act as mediators of *trans* effects. There was extensive colocalisation of causal variants for cell type- and stimulation-specific neonatal *cis*-eQTLs and those of autoimmune and allergic diseases, in particular *CTSH* (Cathepsin H) which showed widespread colocalisation across diseases. Mendelian randomisation showed causal neonatal gene transcription effects on disease risk for *BTN3A2*, *HLA-C* and many other genes. Our study elucidates the genetics of gene expression in neonatal conditions and cell types as well as the aetiological origins of autoimmune and allergic diseases.

## Introduction

Infancy is a critical period during which physiological and developmental changes impact the pathogenesis of conditions later in life^1,2^. Many complex diseases, in particular immune and respiratory conditions, are partially determined by genetic predisposition and early-life environment exposures, such as microbes or allergens^3–5^. Yet, despite increasing evidence of its importance, little is known about early life genetic regulation of gene expression and its relevance to the genetic predisposition for diseases which arise later in life.

Expression quantitative trait loci (eQTL) studies have provided insights into the gene regulatory effects of genetic variants and their relationship with complex disease^6–9^. The majority of eQTLs have been identified in adult tissues, and exploration of eQTLs in perinatal tissues has only recently commenced: for example, eQTLs identified in foetal placentas^10^ and foetal brains^11^ are enriched for genetic variants associated with development (e.g. adult height) and neuropsychiatric (e.g. schizophrenia) traits, respectively. In addition to genetic variation, disease development is influenced by individual response to external stimuli. Understanding the interaction between eQTLs and external stimuli can give insights into the condition(s), whether they be cell type, microbe or temperature, under which genetic variants may exert their effect on disease. Previous studies have investigated response eQTLs (reQTLs), eQTLs with genetic effects modified by external stimulation, in CD14^+^ monocytes^12,13^, macrophages^14^, dendritic cells^15,16^, and CD4^+^ T cells^17^. However, reQTL studies to date have largely been performed using samples from adult tissues and cell types and thus our knowledge of the genetic landscape of neonatal gene expression responses to common immune-mediated stimuli is limited.

Here, we characterise the genetics of gene expression in the innate and adaptive arms of the neonatal immune system using purified cord blood samples from 152 neonates^18–23^. In these samples, we mapped *cis*- and *trans*-eQTLs of monocytes and CD4^+^ T cells as well as reQTLs for monotypes stimulated with lipopolysaccharide (LPS; a component of bacterial cell walls) and CD4+ T cells stimulated with phytohemagglutinin (PHA; a pan-T cell mitogen). Using mediation analysis, putative *trans* gene regulation was investigated to identify *cis* regulatory mechanisms. We explored the shared genetic basis of neonatal eQTLs and reQTLs with common autoimmune and allergic diseases. Finally, we used Mendelian randomisation to uncover causal effects of neonatal monocyte and CD4^+^ T gene expression on disease risk, and to highlight the potential importance of the perinatal period in understanding the origins of immune-mediated disease.

## Results

### Genetics of neonatal gene expression in innate and adaptive immunity

We performed eQTL analysis on *in vitro* cultures of resting and stimulated neonatal immune cells from 152 neonates of the Childhood Asthma Study (CAS) cohort (Figure 1). The cell cultures comprised four experimental conditions: resting monocytes, LPS-stimulated monocytes, resting T cells, and PHA-stimulated T cells. The total number of samples available for eQTL analysis was 116 for resting monocytes, 125 for LPS-stimulated monocytes, 126 for resting T cells, and 127 for PHA-stimulated T cells.

**Figure 1:**
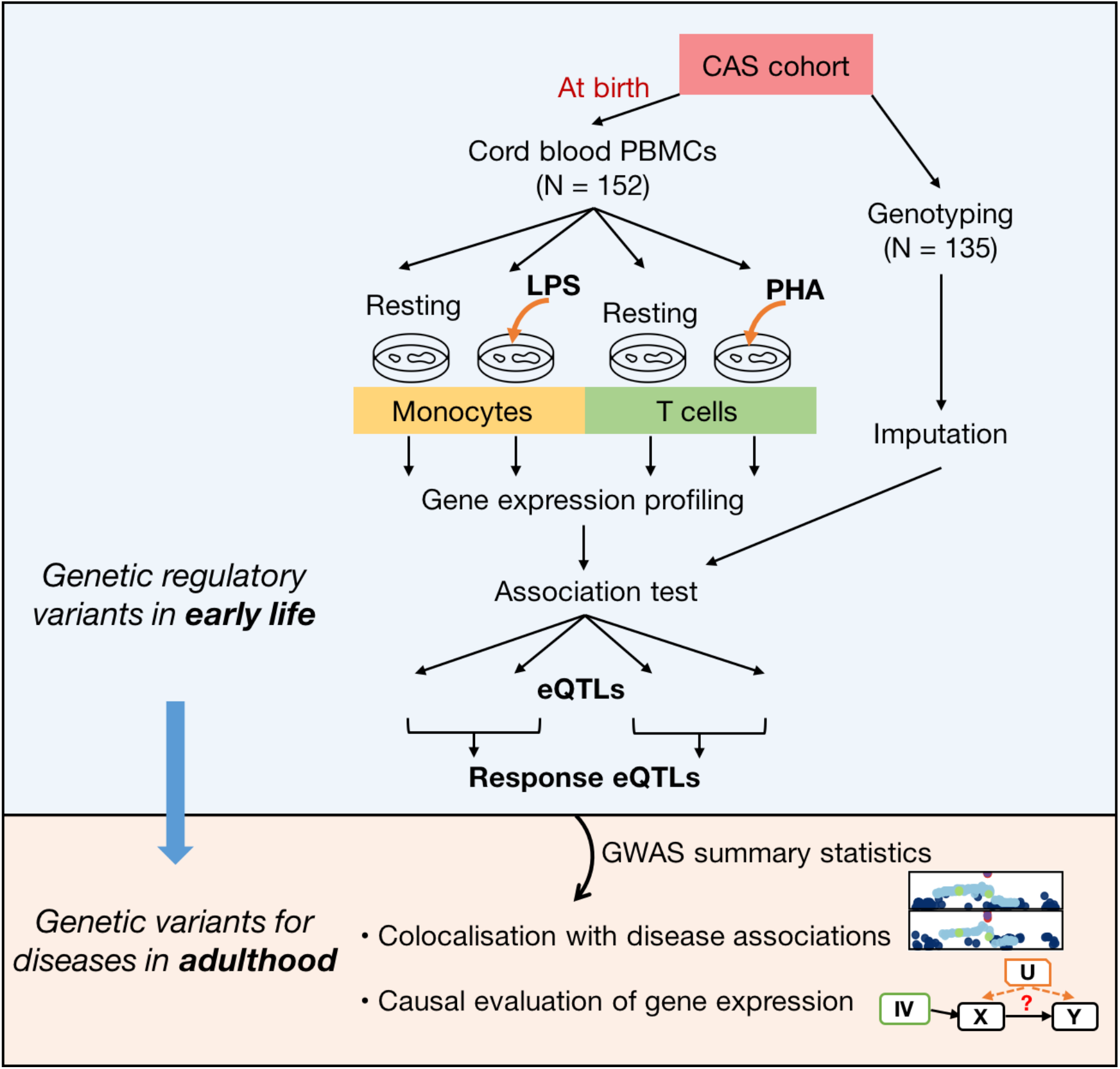
Study design and analysis work flow. Monocytes and T cell cultures were purified from resting and stimulated cord blood samples from the Childhood Asthma Study (CAS) cohort. Gene expression was quantified using a microarray platform. Genotype data are available for a subset of the CAS individuals. eQTLs were identified within each experimental condition. Datasets for resting and stimulated samples were merged to detect response eQTLs within each cell type. Next, we identified genetic loci where neonatal eQTLs and disease associations obtained from external GWAS datasets shared the same causal variants. We investigated the causal effects of gene expression at birth on immune diseases that develop later in life.

To identify *cis-*eQTLs, we applied a hierarchical procedure to correct for multiple testing within each experimental condition at 5% false discovery rate (FDR; **Methods**). Stimulated cells yielded a larger number of *cis-*eQTLs and associated genes (eGenes) than resting cells (1,347 vs. 971 eGenes in PHA-stimulated vs. resting T cells, respectively; 376 vs. 136 in LPS-stimulated vs. resting monocytes, respectively; Figure 2A, **Table S1–4**). To investigate the differences in numbers of eGenes between conditions, we repeated the analysis controlling for differences in sample size (randomly sampling 116 samples in each condition). This yielded similar results to the numerical distribution of *cis-*eGenes: 1,231, 900, and 350 in PHA-stimulated T cells, resting T cells, and LPS-stimulated monocytes, respectively. The lower number of eQTLs in monocytes may be explained by fewer genes being expressed (**Figure S1**).

**Figure 2:**
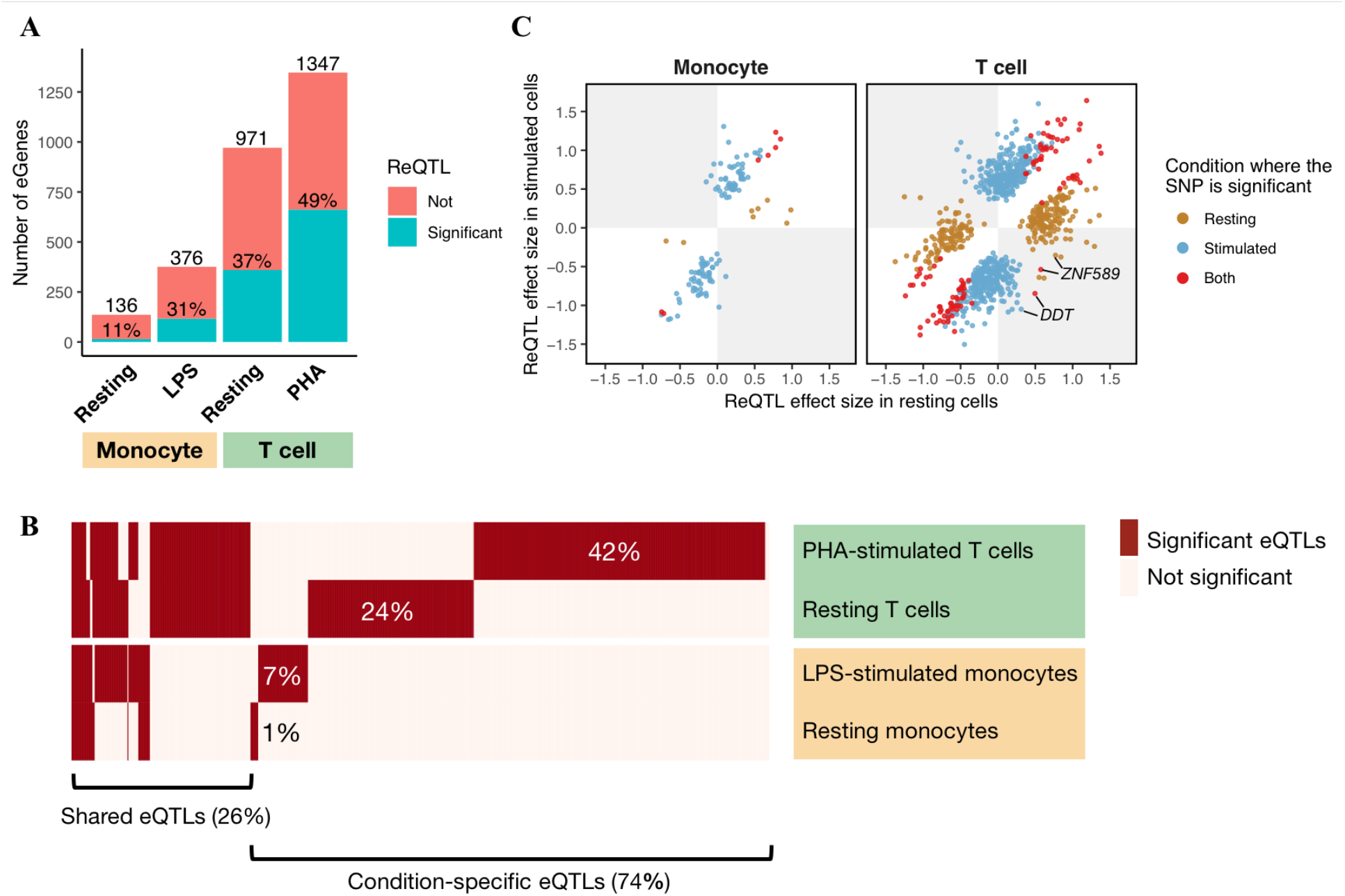
*Cis*-eQTLs and response eQTLs (reQTLs) in monocytes and T cells. (**A**) A bar plot shows the number of genes with significant *cis*-eQTLs (eGenes) identified in each cell type and treatment group (on x-axis). Percentages on each bar indicate the proportion of eGenes with significant reQTLs (green). (**B**) A heatmap shows eQTL sharing across four experimental conditions (rows). Columns in the heatmap are unique eQTL–eGene associations. Significant associations are in red. Percentages labelled on the heatmap show the proportion of unique eQTL–eGene associations that are specific to a certain cell type and stimulatory condition. (**C**) Two point plots show effect sizes (s.d. change in gene expression per allele) of significant reQTLs in resting (x-axes) and stimulated conditions (y-axes) in two cell populations: monocytes (left) and T cells (right). A gene might have two dots indicating two independent top SNPs (**Methods**). Colours indicate the condition in which the SNP was significant. ReQTLs of *DDT* and *ZNF585* in the grey quadrants (red dots) show opposite directions of eQTL effects across conditions.

For eGenes with eQTLs in multiple experimental conditions, we performed conditional analysis to distinguish whether these were independent or shared signals between conditions (**Methods**). The majority (74%) of eQTL signals were specific to one cell type or stimulatory condition (Figure 2B), consistent with previous observations^12^. We observed a majority of *cis*-eQTL effects after stimulation: 60% (262 of 376) of eGenes in LPS-stimulated monocytes and 58% (778 of 1,347) in PHA-stimulated T cells. Using a two-step conditional analysis (**Methods**), PHA-stimulated T cells had the largest number of eGenes (6.3%; **Table S5**) with multiple independent eQTL signals. *GARFIELD* enrichment analysis^24^ showed that the *cis*-eSNPs were enriched in 3’ untranslated regions (UTR), 5’ UTR, and exon regions (**Figure S2**), consistent with known mechanisms of *cis-*eQTLs.

In comparing our resting and LPS-stimulated monocytes to those from adults in Fairfax *et al.*^12^, we found that approximately half of the *cis*-eQTLs from neonatal monocytes (including variants in high LD, r^2^ ≥0.8) replicated in adult monocytes (**Table S6**). Similarly, we compared resting and stimulated neonatal T cells to adults T cells of the DICE study^7^: although the stimuli were different, 32% and 18% of *cis*-eQTLs from resting and PHA-stimulated CD4^+^ T cells in neonates replicated in resting and anti-CD3/CD28-stimulated CD4^+^ T cells, respectively.

### Genetics of neonatal gene expression in response to stimuli

To quantify how genetic regulation of gene expression is altered by external stimuli, we identified response eQTLs (reQTLs) and response eGenes (reGenes) by performing interaction tests on the top eSNPs of each eGene in monocytes and T cells separately, and controlling FDR at 5% using permutation-adjusted P-values (**Methods**). In monocytes, we identified 125 significant reQTLs involving 125 unique reGenes (31% of 398 monocyte eGenes); in T cells, we identified 956 reQTLs involving 918 unique reGenes (52% of 1,749 T cell eGenes), among which 38 reGenes had distinct *cis*-eQTLs in two conditions where both eQTLs were reQTLs (**Table S7–8**). Consistent with our findings for *cis* eQTLs and eGenes, the number of reQTLs and proportion of reGenes were greater in stimulated compared to resting conditions.

For two reQTLs, the direction of eQTL effect changed between conditions (Figure 2C). The ‘C’ allele of the top eSNP (rs5751775) for *DDT* (D-dopachrome tautomerase), a gene functionally related to the inflammatory cytokine *MIF* (migration inhibitory factor), increased *DDT* transcription in resting T cells but decreased expression after PHA stimulation (**Figure S3A**, **Table S8**). Similarly, the ‘T’ allele of the top eSNP (rs13068288) for *ZNF589* increased transcription of *ZNF589* in resting T cells but reduced expression after PHA stimulation (**Figure S3B**, **Table S8**).

### Disentangling *trans* and *cis* effects using mediation analysis

We identified 25 *trans*-eQTLs in T cells (10 in resting, 15 in PHA-stimulated), and one *trans*-eQTL in monocytes (Figure 3A, **Table S9**) at a genome-wide FDR of 5%. Notably, the *trans*-eQTL for *MYH10*, a component of myosin heavy chain which regulates cytokinesis, was shared across all four experimental conditions; furthermore, the same eQTL was associated with multiple *trans*-eGenes in T cells: *MIR130A* and *STX1B* in resting and stimulated T cells, and *IP6K2* and *MIR1471* in resting T cells only (Figure 3A).

**Figure 3:**
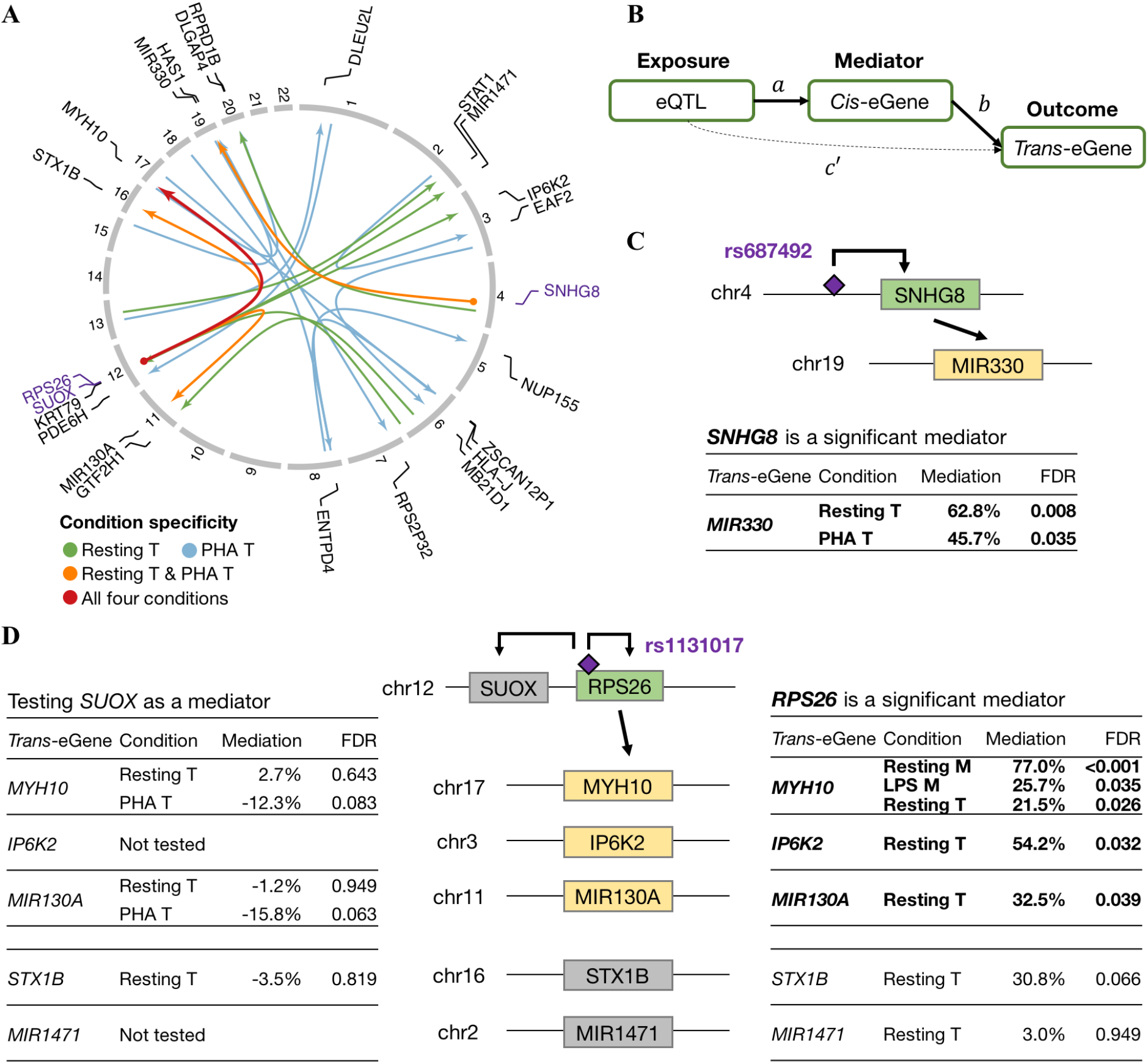
*Trans*-eQTL effects and their *cis-*mediators. (**A**) A circular plot shows *trans*-eQTL associations in lines, with arrows pointing to *trans*-eGenes with names annotated (in black) outside the rim indicating chromosome numbers. Dots on the other end point indicate nearby genes (names in purple) that are associated with the same loci (*cis*-eQTLs). Colours of the lines indicate the experimental conditions where the *trans*-eQTLs were identified: “Resting T”: resting T cells only, “PHA T”: stimulated T cells only, “Resting T & PHA T”: shared between both conditions of T cells, and “All four conditions”: shared across all four experimental conditions. (**B**) A diagram demonstrates the mediation analysis model, where effects of *trans*-acting eQTL (exposure) on *trans-*eGene (outcome) are either mediated through a *cis-*eGene (mediator), or through direct effects (**Methods**). (**C**) and (**D**) show two examples of *cis*-eGenes (green), *SNHG8* and *RPS26*, acting as mediators for *trans*-effects (*trans*-eGenes in yellow). Genes that were not significant in mediation analysis are in grey. Tables show statistics of the mediation tests, and the “Mediation” column indicates the proportion of total effects of the eQTL on the *trans*-eGene that was mediated through the *cis*-eGene. Two models involving *SUOX* (**D**) were not tested because the *trans*-eSNPs of *IP6K2* and *MIR1471* were not significantly associated with *SUOX* (**Table S9**). Significant mediations (FDR ≤0.05) are highlighted in bold.

Consistent with previous reports that *trans*-eQTLs are enriched for *cis*-eQTLs^25^, we found that multiple *trans*-eQTLs (the single *trans*-eQTL in monocytes, 6 of the 10 in resting T cells, and 3 of the 15 in PHA-stimulated T cells) were significantly associated with local genes in *cis* (Figure 3A, **Table S9**). The *trans*-eQTL for *MYH10* was also a *cis*-eQTL for *RPS26* (part of the 40S subunit of the ribosome) in all conditions except PHA-stimulated T cells. The top eSNP (rs1131017) for *RPS26*, located in its 5’ UTR, was also a *cis*-eQTL for *SUOX* (sulphite oxidase, a homodimer in the intermembrane space of mitochondria) in resting and stimulated T cells. Separately, the *trans*-eQTL (rs687492) for microRNA *MIR330* was also a *cis*-eQTL for the long non-coding RNA *SNHG8* in resting and stimulated T cells.

Mediation analysis revealed that the *trans*-eQTL effects of rs687492 on *MIR330* were *cis* mediated through *SNHG8* (Figure 3B–C, **Table S10**, **Methods**), indicating a potential pathway containing this lncRNA-miRNA cross-talk. Furthermore, mediation analysis also revealed the regulatory logic of the *cis*-eQTL (rs1131017) for *SUOX* and *RPS26*, revealing that rs1131017’s *trans-* effects on *MYH10, MIR130A, IP6K2* were mediated through *RPS26* and not *SUOX* in resting T cells (Figure 3D, **Table S10**).

### Genetic overlap with immune-mediated diseases

To investigate the genetic overlap between neonatal gene expression and disease, we used a multi-pronged approach. First, we performed enrichment analyses to test for significant overlaps between the *cis*-eQTLs and variants associated with immune-mediated disease in genome-wide association studies (**Methods**). We found widespread enrichment amongst *cis*-eQTLs for genetic variants associated with diseases such as allergic disease (asthma, hay fever, or eczema) and inflammatory bowel disease (**Figure S4**).

Second, we performed colocalisation analysis^26^ to identify variants sharing regulatory (eQTL) and disease-associated (GWAS) signals (**Methods**). In total, we observed 68 colocalisations, involving 5, 9, 15, and 17 independent *cis-*eQTLs in resting monocytes, LPS-stimulated monocytes, resting T cells, and PHA-stimulated T cells, respectively (Figure 4A, **Table S11**). Our analysis revealed widespread colocalisation of the *cis*-eQTL for *BACH2* in resting T cells with variants for autoimmune thyroid disease, celiac disease, multiple sclerosis, rheumatoid arthritis, and type 1 diabetes (**Figure S5**). *BACH2* encodes a transcriptional repressor that restrains terminal differentiation and promotes the development of memory lymphocytes including CD8^+^ T cells^27^ and B cells^28^. At the *BACH2* locus, the ‘A’ allele at the top eSNP (rs72928038) was associated with decreased *BACH2* expression and increased risk of the above diseases. This was consistent with previous studies which showed mutations and loss-of-function variants of *BACH2* resulted in immunodeficiency and disruption to regulatory T cells function with subsequent autoimmunity^29,30^.

**Figure 4:**
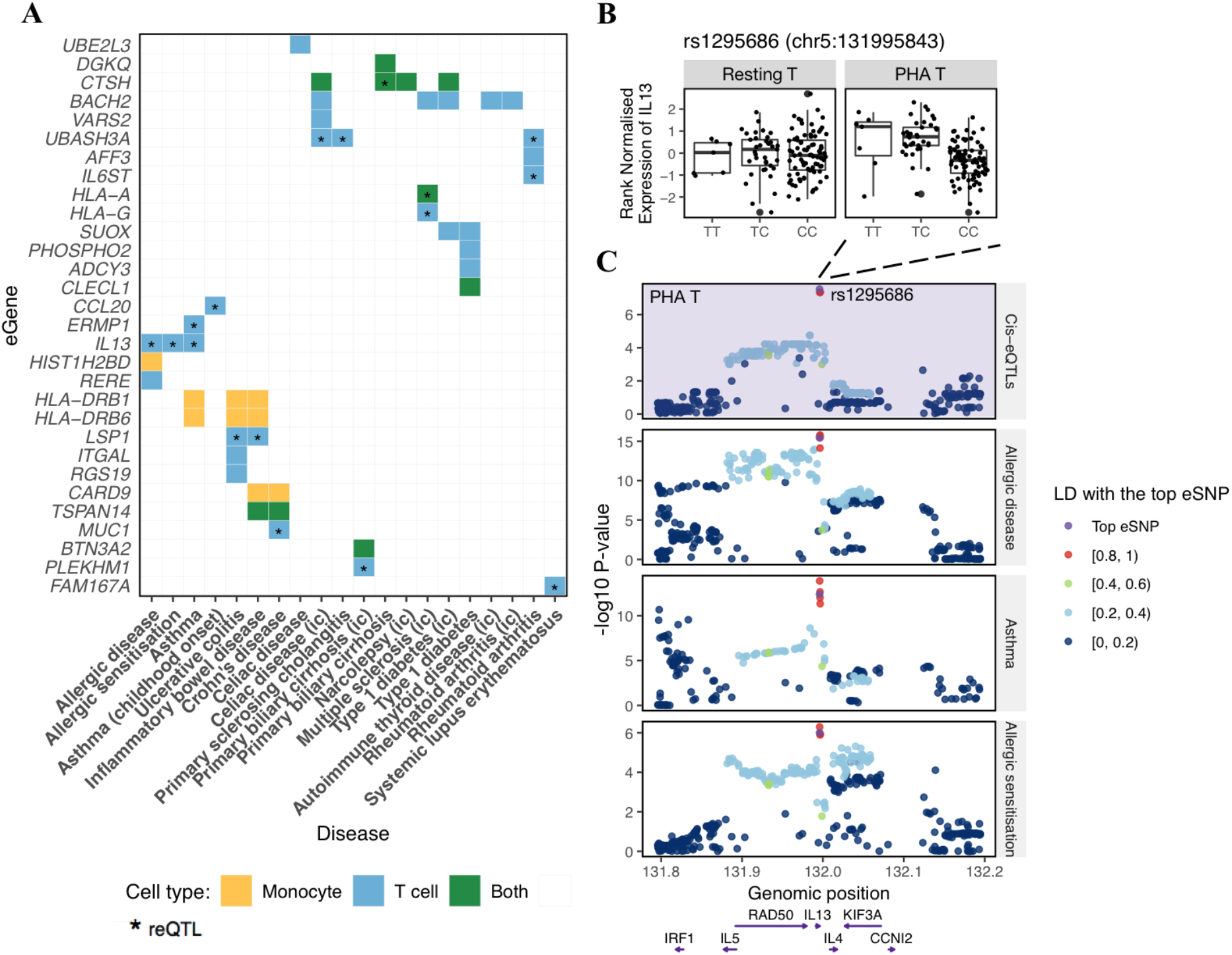
Colocalisation of *cis*-eQTLs with disease associations. (**A**) A heatmap shows all cases with strong evidence of colocalisation between *cis-*eQTLs of corresponding genes (eGenes) in rows and GWAS hits associated with allergic and autoimmune diseases in columns (“ic” indicates that the study was performed using ImmunoChip array). Colours indicate the cell type where the significant colocalisation was observed. Asterisks indicate that the colocalised eQTLs are response eQTLs (reQTLs). (**B**) Box plots show the rank-normalised gene expression of *IL13* (y-axes) in resting T cells (left) and in PHA-stimulated T cells (right) stratified by genotypes of the reQTL rs1295686 (x-axes), the top eSNP in PHA-stimulated T cells. In resting T cells, no SNP was significantly associated with *IL13*. (**C**) Regional plots show eQTL association with gene expression of *IL13* in PHA-stimulated T cells (purple background), and GWAS associations with allergic disease (asthma, hay fever, or eczema), asthma, and allergic sensitisation. The minus log10 P-value is plotted on y-axes for all SNPs located within 200 kb from the top eSNP of *IL13*. Colours of dots indicate the LD correlation with the top eSNP (in purple). Positions of genes located on this locus are shown at the bottom.

We found 17 colocalisations of reQTLs and disease variants, in total involving 12 reQTLs: one monocyte reQTL specific to LPS stimulation (eQTL for *CTSH*), and 11 reQTLs in T cells, among which 8 were specific to PHA stimulation. Notably, the reQTL for *IL13* in PHA-stimulated T cells colocalised with GWAS hits associated with asthma, allergic sensitisation, and allergic disease (Figure 4, **Table S11**). The ‘T’ allele of the top eSNP (rs1295686) was associated with greater *IL13* expression in PHA-stimulated T cells as well as increased risk of all three diseases (Figure 4B). rs1295686 is intronic to *IL13* and in strong LD (r^2^ >0.98) with four other eSNPs, including an Gln144Arg missense SNP (rs20541) in *IL13*. At the *CCL20* locus, the ‘A’ allele of the *cis*-eQTL/reQTL (rs13034664) in PHA-stimulated T cells was associated with lower *CCL20* expression as well as increased risk of childhood-onset asthma (Figure 4A, **Figure S6**, **Table S11**). *CCL20* is part of the *CCR6*-*CCL20* receptor-ligand axis, a key driver of dendritic cell chemotaxis^31^.

Our analyses uncovered complex condition-specific colocalisations at multiple loci. A T cell ubiquitin ligand that regulates apoptosis (*UBASH3A*) had two independent *cis*-eQTLs in resting and stimulated T cells, which were also reQTLs. However, only the *cis*-eQTL in resting T cells (rs1893592) colocalised with celiac disease, rheumatoid arthritis, and primary sclerosing cholangitis (PSC) (Figure 4, **Table S11**). *Cis*-eQTLs for *CTSH*, which encodes the lysosomal cysteine proteinase cathepsin H, colocalised with signals for different diseases in a cell type and condition specific manner (**Figure S7**, **Table S11**). *Cis-*eQTLs for *CTSH* in resting monocytes and resting T cells both colocalised with GWAS hits for celiac disease, narcolepsy, and type 1 diabetes. On the other hand, *cis-*eQTLs for *CTSH* in LPS-stimulated monocytes and PHA-stimulated T cells both colocalised with primary biliary cirrhosis (PBC).

### Causal effects of condition-specific gene expression on immune-mediated diseases

To identify putative causal effects of neonatal gene expression on risk of autoimmune and allergic disease, we performed two-sample Mendelian randomisation (MR) analysis using *cis*-eQTLs as genetic instruments, the neonatal *cis*-eGene as exposure, and disease as outcome (**Methods**). We tested the 52 eGenes which had 3 or more genetic instruments available and the diseases above, for which we had GWAS summary statistics available. We considered genes for which at least three of four MR methods (inverse variance weighted, weighted median, weighted mode, and MR Egger) were in agreement in detecting significant causal effects (P-value ≤0.05) on a disease without significant pleiotropic effects (**Table S12**).

In our MR analysis, we found multiple conditions where neonatal gene expression had a causal effect on multiple diseases (Figure 5), including *BTN3A2* (butyrophilin subfamily 3 member A2), *HLA-C* (major histocompatibility complex class I molecule), *MICB* (ligand for an activatory receptor expressed on natural killer cells, CD8^+^ αβ T cells, and γδ T cells), *ZNRD1* (RNA polymerase 1 subunit), and *SLC22A5* (carnitine transporter) (**Table S12**). *BTN3A2* had a relatively large number of genetic instruments for resting (7 to 8) and stimulated (3 to 5) T cells, and the causal estimates were similar between these two conditions (**Figure S8–9)**. In resting T cells, increased expression of *BTN3A2* was causally associated with decreased risk of asthma (weighted mode causal estimate = −0.056 log odds decrease per s.d. increase in *BTN3A2*), both childhood-and adult-onset asthma (−0.047 and −0.039, respectively), allergic rhinitis (−0.044), PSC (−0.440), and systemic lupus erythematosus (SLE; −0.256). Conversely, increased *BTN3A2* expression was associated with increased risk of inflammatory bowel disease (IBD; 0.025), including Crohn’s disease (0.053), as well as risk of PBC (0.129), where PBC variants also showed colocalisation with *BTN3A2* eQTLs (Figure 4, **Figure S10**). Expression of *HLA-C* in T cells showed strong causal association with autoimmunity, in particular positive causal effects on psoriasis, SLE, PSC, multiple sclerosis, IBD, and ulcerative colitis; and negative causal effects on juvenile idiopathic arthritis, PBC, and rheumatoid arthritis (Figure 5).

**Figure 5:**
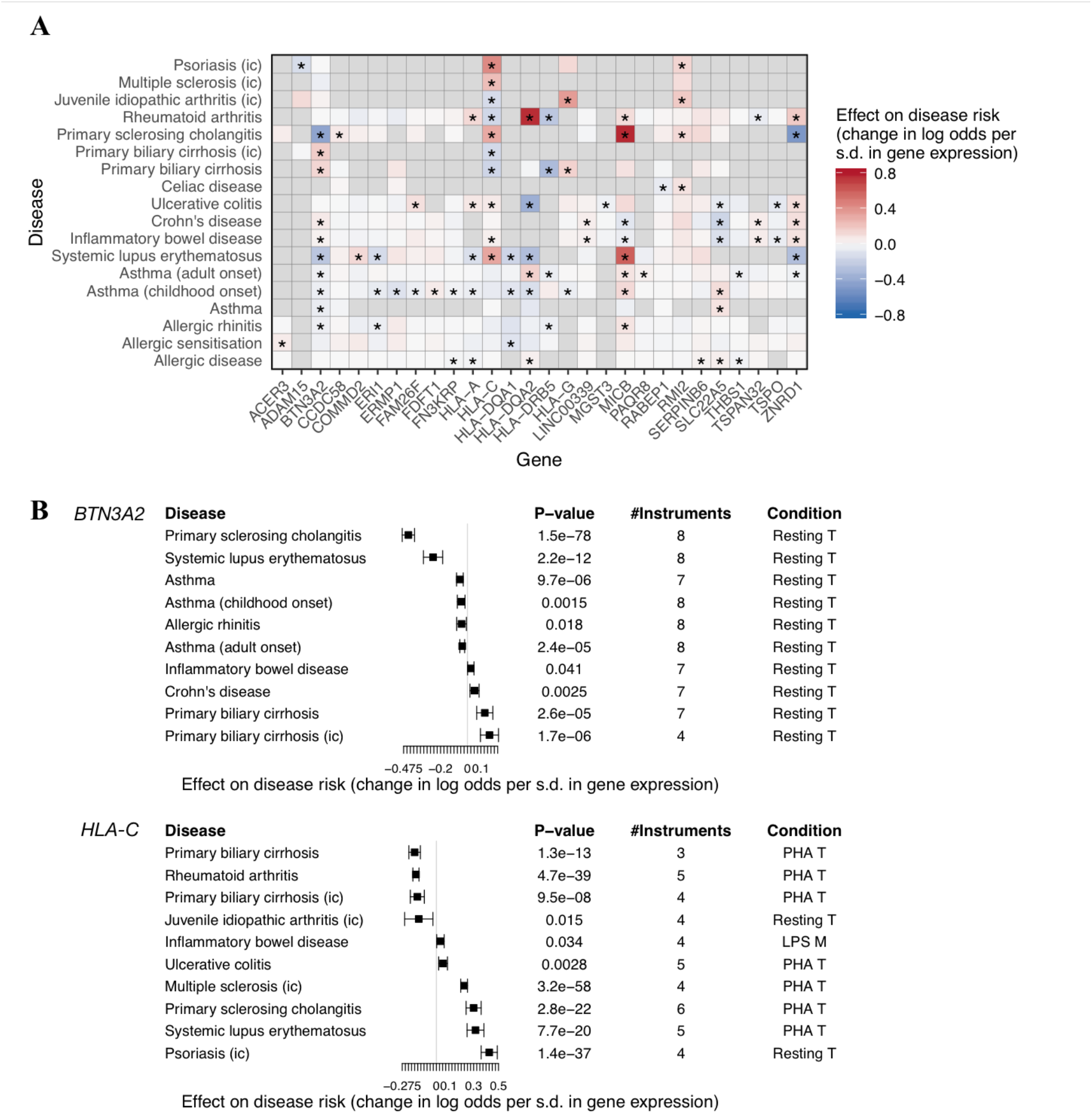
Causal effects of neonatal gene expression on multiple immune-related diseases. (**A**) The heatmap shows the causal effect size estimated using the weighted mode method in the Mendelian randomisation (MR) analysis. Asterisks indicate significant causal associations (**Methods**). Causal associations with significant pleiotropy was excluded. If a gene was tested using the expression levels in multiple experimental conditions, the one with the highest number of genetic instruments was kept. Statistics of all MR tests are in **Table S12**. Grey indicates the gene-disease pairs that were not tested due to small number of genetic instruments (<3). Positive effect estimates in red indicate that increased gene expression is causally associated with increased disease risk, and negative causal associations are in blue. (**B**) Forest plots show causal estimates and 95% confidence intervals for neonatal expression of *BTN3A2* and *HLA-C*.

## Discussion

In this study we investigated the genetic regulation of gene expression in the innate and adaptive arms of the neonatal immune system and its relationship to the genetic basis of autoimmune and allergic diseases. In this context, we illustrated that the genetics of gene expression in neonates is strongly specific to cell type and stimulatory condition. We described regulatory mechanisms of eQTLs whose *trans* effects are mediated via gene expression in *cis*. In exploring the potential early life origins of disease, our analyses showed extensive genetic overlap of genetic variants associated with immune-mediated diseases and those with effects on gene expression in neonatal immune cells. Finally, Mendelian randomisation showed that myriad changes in gene expression at birth had potentially causal effects on autoimmune and allergic disease risk.

We observed stimuli changing the direction of eQTL effects in resting and PHA-stimulated T cells. This was the case for the ‘C’ allele of reQTL (rs5751775) and *DDT*, a cytokine structurally and functionally related to *MIF*, a critical regulator of both innate and the adaptive immune response^32,33^. It is known that eQTL effects can change direction in immune cells^34^. In the Genotype-Tissue Expression (GTEx) Project^6^, the direction of eQTL effect for the ‘C’ allele at rs5751775 was also variable across liver, pancreas, stomach, testis, brain, and muscle tissues.

Our results revealed a *trans* mediation role for *RPS26*, rather than *SUOX*, consistent with previous studies linking *RPS26* to *IP6K2*^35,36^. *RPS26* and *SUOX* were both identified as significant mediators in multiple tissues in the GTEx dataset; however, within the same tissue, the *trans*-associations mediated through *SUOX* were not identical to those through *RPS26*, indicating distinct effects of *SUOX* and *RPS26* on distant genes^36^. *RPS26* encodes a ribosomal subunit protein which, apart from its role in ribosome assembly and translation^37,38^, is involved in various other cellular processes, including nonsense-mediated mRNA decay^39^ and p53 transcriptional activity^40^. It is likely that the broad *trans* effects of *RPS26* are related to its ribosomal functions.

At the *CCL20* locus, the reQTL (rs13034664) in PHA-stimulated T cells colocalised with childhood-onset asthma^41^ (Figure 4A, **Figure S6**). *CCL20* encodes a C-C chemokine ligand that binds to a G protein-coupled receptor, and elevated CCL20 expression has been shown in airways of patients with chronic obstructive pulmonary disease (COPD)^42^ and asthma^43,44^. CCL20 induces mucin production by binding to its unique receptor (CCR6) in human airway epithelial cells^45^. However, in PHA-stimulated T cells, the ‘A’ allele of the reQTL (rs13034664), which was linked to increased risk of childhood-onset asthma, was associated with reduced *CCL20* expression (**Figure S6B**). This reQTL and its direction of effect were replicated by others in activated CD4^+^ T cells^7^. T cells themselves respond to CCL20 via binding to CCR6, and this process is observed during allergen provocation^46^.

On the other hand, the reQTL for *IL13* in neonatal PHA-stimulated T cells appeared to share causal variants with allergic disease and asthma, suggesting that this reQTL may affect allergic disease risk via mechanisms involving T cell activation and interleukin 13 (IL-13). PHA is a pan T cell mitogen^47^, and downstream intracellular signalling may be shared between PHA and allergen-mediated T cell activation. IL-13 is produced by activated CD4^+^ and CD8^+^ T cells^48^ among others, promoting immunoglobulin E (IgE) production in B cells^49^. IL-13 has been shown to induce asthma symptoms including airway hyper-responsiveness, increased total serum IgE, and increased mucus production in murine models^50^. Increased IL-13 expression is observed in sputum and bronchial biopsy in mild^51^ and severe^52^ asthma and can serve as a biomarker for severe refractory asthma^53^. Therapies that target IL-13 (anti-IL-13 antibodies) have been developed, such as lebrikizumab^54^ and tralokinumab^55^; however, they show inconsistent or only modest effects in treating severe asthma exacerbations in phase 3 clinical trials. Our results are consistent with IL-13 as a therapeutic target for asthma and may suggest potentially increased efficacy in genetic subgroups.

We found strong causal effects of neonatal *BTN3A2* expression on various autoimmune and allergic diseases. Butyrophilin (BTN) family members are immunoglobulin-like molecules that act as immune check-point regulators with roles in self-tolerance^56^. Increased BTN3A2 protein expression is a favourable prognostic biomarker in epithelial ovarian cancer patients, and indicates a higher density of intraepithelial infiltration of T cells^57^. BTN3 family members *BTN3A1* and *BTN3A3* are proximal to *BTN3A2*. The antigen-presenting BTN3A1 is critical to human γδ T cell activation^58,59^. A recent study showed that BTN3A2 regulated subcellular localisation of BTN3A1, and both were required for T cell activation^60^. Previous MR analysis found *BTN3A2* lung expression had a causal effect on COPD risk^61^. Our findings suggest that altered neonatal *BTN3A2* expression, with presumed subsequent dysfunctional immunomodulation, plays a role in the pathogenesis of multiple inflammatory conditions.

In our study, neonatal *HLA-C* expression in both monocytes and T cells was causally associated with multiple autoimmune diseases such as psoriasis, SLE, and primary biliary cirrhosis. *HLA-C* encodes an MHC Class I receptor which presents antigens to CD8+ T cells. It is also the major ligand for killer immunoglobulin-like receptors (KIRs), which regulate the activity of natural killer (NK) cells. *HLA-C* is an established locus for psoriasis susceptibility^62^ and the interaction between *HLA-C* and *ERAP1* is associated with psoriasis risk, where *ERAP1* variants only have psoriasis effects in individuals with the *HLA-C* risk allele^63^. In our analysis of resting neonatal T cells, the largest causal effect of *HLA-C* expression was for psoriasis.

In conclusion, our study shows the remarkable complexity of the genetic regulation of gene expression in the innate and adaptive arms of the immune system at birth, and its potential role in the pathogenesis of autoimmunity and allergic disease.

## Methods

### Study cohort and RNA sample preparation

The study population is a subset of the Childhood Asthma Study (CAS), a prospective birth cohort of 234 individuals followed from birth to up to 10 years of age^18–23^. Cord blood samples were collected for 152 individuals at birth. One million peripheral blood mononuclear cells (PBMCs) from each individual were stimulated with either an innate immune system stimulant (LPS: lipopolysaccharide), or a pan T cell stimulant (PHA: phytohemagglutinin) for 24 hours (Figure 1). Unstimulated resting PBMC samples were also available. Non-adherent cells in suspension from resting and PHA-stimulated cultures were removed and purified for CD8^−^/CD4^+^ T cells using Dynabeads (Invitrogen) and stored in RNAprotect Cell reagent (Qiagen). Cells remaining in suspension in resting and LPS-stimulated cultures were aspirated, leaving an enriched population of monocytes and macrophages adhered to the culture wells. These adherent cells were then resuspended and transferred into RNAprotect. All cells in RNAprotect Cell reagent were banked at −80°C.

The cells were thawed and centrifuged briefly for RNA extraction. Reagent was removed and total RNA was extracted from pelleted cells by an established in-house procedure using TRIzol (Life Technologies) in combination with RNEasy MinElute columns (Qiagen). The aqueous phase containing the RNA was then loaded onto an RNeasy MinElute column (Qiagen) to purify and concentrate the RNA. RNA quality was assessed on a Bioanalyzer 2100 using the RNA 6000 Nano kit (Agilent). There were 607 samples (one missing sample) in total from 152 individuals for gene expression profiling.

### Gene expression profiling and data processing

Total RNA from four cell culture conditions (resting and LPS-stimulated monocytes, and resting and PHA-stimulated T cells) was quantified with Illumina HumanHT-12 v4 BeadChip gene expression array at the Genome Institute of Singapore. After excluding 31 samples with suspected cross-contamination or insufficient quantity of cDNA, 576 samples were successfully scanned. The raw microarray data and probe detection P-values were exported by the Illumina software *BeadStudio*. We first removed 3 samples with zero intensity for almost all probes including negative controls and probes targeting housekeeping genes (2 resting monocyte samples and 1 LPS-stimulated monocyte sample). We further removed 16 outlier samples (8 resting monocyte samples, 5 LPS-stimulated monocyte samples, and 3 resting T cell samples) with a low number of detectable probes (lying outside median ±2 × inter quartile range). Compared with other samples, these excluded samples had much lower intensity for positive controls including those targeting housekeeping genes.

After quality control, 557 samples remained for normalisation (resting monocytes: 130, LPS-stimulated monocytes: 141, resting T cells: 142, and PHA-stimulated T cells: 144). We performed background correction based on the intensity of the 770 negative control probes on the microarray, and then we performed quantile normalisation and log2 transformation within each cell type and condition using the *neqc* function from the *limma* R package^64^. We used updated probe annotation data and restricted the analysis to 33,436 reliable probes, excluding unaligned probes and probes aligned to multiple regions that were more than 25 bp apart^65^. Fifteen probes with missing data in ≥5 samples were removed. Detectable probes targeting autosomal genes (N = 20,532) were kept, comprising of probes with *BeadStudio* detection P-values ≤0.01 in ≥2.5% of the samples from a specific condition group, or in ≥5% of all samples^12^. Gene annotation was obtained from the GENCODE^66^ release 19 (GRCh37 alignment, downloaded in October 2017). Among detectable probes, 19,230 had gene annotation in the GENCODE reference data. For genes that had multiple probes, we kept the probe with the highest mean intensity^67^, resulting in 13,109 autosomal genes. For eQTL analysis, we performed a rank-based inverse normal transformation within each group, so that each gene expression followed a standard normal distribution.

### Genotyping and imputation

Genomic DNA was extracted from blood samples collected from 218 individuals. Genotyping was performed with Illumina Omni2.5 BeadChip array, with coverage of approximately 2.5 million markers. Variants with missing call rates >1%, MAF <1%, or Hardy-Weinberg Equilibrium (HWE) test P-value <1×10^−6^ were excluded, and individuals with missing call rates >1% were removed. This produced an initial count of 1.4 million SNPs for 215 genotyped individuals. Of these, a total of 135 children also had gene expression data from cord blood (i.e. overlap with the 152 individuals described previously).

We performed genotype imputation using the Michigan Imputation Server^68^ with Haplotype Reference Consortium^69^ (HRC) release r1.1 as the reference panel. After filtering out variants with low imputation accuracy (R^2^ <0.3), 12.7 million SNPs remained. For eQTL analysis, we focused on 4.3 million SNPs with MAFs ≥10%. The MAF cut-off used here was suggested by an eQTL simulation study in order to avoid inflated false positives in low-frequency variants given our limited sample size^70^.

### *Cis*-eQTL mapping and conditional analysis

To identify *cis-*eQTLs within each cell type and treatment group, we performed linear additive regression to model the effect of each SNP located within 1Mb of the transcription start site (TSS) of the corresponding gene using the *Matrix eQTL* R package^71^. The sample size for eQTL mapping in each experimental condition was: 116 for resting monocytes, 125 for LPS-stimulated monocytes, 126 for resting T cells, and 127 for PHA-stimulated T cells. Genotype data were recoded as 0, 1, 2 based on the dosage of the HRC alternative allele. Gender, first three genotype PCs^72^ and first ten PEER^73^ factors capturing technical variation in transcriptomes were included as covariates in the linear model.

We applied a hierarchical correction procedure to correct for multiple testing^70^. Firstly, nominal P-values for all *cis*-SNPs from *Matrix eQTL* were adjusted by multiplying the number of effective independent SNPs for each gene (local correction), which was estimated by *eigenMT* based on genotype correlation matrix^74^. Secondly, the minimum locally adjusted P-value for each gene was kept and the FDR of significant genes was controlled at 5% using the Benjamini-Hochberg (BH) FDR-controlling procedure (global correction)^75^. Genes with global FDR ≤0.05 were considered significant eGenes. Thirdly, to obtain the list of significant eSNPs for each eGene, the locally adjusted minimum P-value corresponding to the global FDR threshold of 0.05 was calculated, and SNPs with a locally adjusted P-value lower than the threshold were considered significant eSNPs.

Next, we performed conditional analyses to identify additional independent eQTL signals for each eGene. The gene-level P-value nominal thresholds calculated in the hierarchical multiple-testing correction (eigenMT-BH) were used to determine significant associations: the locally adjusted minimum P-value corresponding to the global FDR threshold of 0.05 multiplied by the number of estimated independent SNPs for each gene. We used a two-step conditional analysis scheme as follows^76^:

#### (1) Forward stage

For each eGene, the number of independent *cis*-eQTL signals was learnt from an iterative procedure. We started from the top SNP with the minimum P-value for the eGene, which was added as a covariate in the linear model to test for *cis*-eQTLs. If any significant SNPs (with P-values smaller than the gene’s nominal threshold) were identified, the new top SNP identified in this iteration was added to the list of independent eQTL signals. In the next iteration of eQTL mapping, all previously identified eSNPs were adjusted for as covariates. The forward stage terminated if no additional significant associations were identified.

#### (2) Backward stage

In this stage, the final list of significant SNPs representing each independent eQTL signal was determined. Let the list of independent SNPs for each eGene obtained from the forward stage be *SNP*_1_, …,*SNP*_2_,*SNP*_3_, …, *SNP*_*M*_, where *M* is the number of independent eQTL signals. Each of the independent eQTL signals was tested separately using a leave-out-one model adjusting for all other SNPs in the list as covariates. For example, when the *i*^th^ eQTL signal was tested, *SNP*_1_, …, *SNP*_*i*−1_, *SNP*_*i*+1_, …, *SNP*_*M*_) were added as covariates together with other covariates used in the original eQTL scan. The final set of independent eQTLs comprised of the eSNPs that remained significant in the backward stage.

For genes that had eQTLs in more than one experimental condition, we also applied conditional analyses to identify independent eQTL signals between conditions. More specifically, to determine whether two eQTL signals for an eGene identified in two experimental conditions were independent or the same signal, we adjusted for the top eSNP in one condition by adding it as a covariate in the linear model and performed eQTL scan again in the other condition. If any SNP was significant in the conditional model using the P-value threshold determined by the hierarchical correction procedure (eigenMT-BH), we considered these two eQTLs as independent signals. If none were significant in the conditional model, we considered it as shared eQTL signal between two conditions or lack of power to detect the independent signal.

### Replication of *cis*-eQTLs in external datasets

We downloaded summary statistics of significant *cis*-eQTLs from a response eQTL study (Supplemental Table S2 of the Fairfax *et al*. study^12^) and the DICE (database of immune cell expression, eQTLs, and epigenomics) project (**URLs**)^7^. Fairfax *et al.* mapped *cis*-eQTLs in resting and LPS-stimulated CD14^+^ monocytes (with two different durations of LPS: 2 or 24 hours) obtained from adults aged from 19 to 56 years, with the sample size being 414 for resting monocytes, and 261 and 322 for monocytes treated with LPS for 2 hours and 24 hours, respectively^12^. The downloaded *cis-*eQTLs were significant at FDR 5% using pooled BH FDR method (i.e. BH FDR-controlling procedure applied to all tests). In the DICE study, 13 immune cell types were collected from 91 subjects, and CD4^+^ T cells and CD8^+^ T cells also had activated conditions^7^. We downloaded eQTLs (P-value ≤1×10^−4^) identified in CD14^+^ monocytes, CD4^+^ T cells, and activated CD4^+^ T cells (4-hour treatment with anti-CD3/CD28 antibodies). In each of the four experimental conditions, we focused on top eSNPs and eSNPs in high LD (r^2^ ≥0.8) for each eGene, and if any of these eSNPs were significantly associated with the same eGene in the corresponding conditions in the Fairfax *et al.* and the DICE datasets, this eQTL signal was considered as replicated.

### Enrichment analysis

We performed enrichment analyses using *GARFIELD* (version 2) to investigate the enrichment patterns of *cis*-eQTLs using predefined features (“annotation data”) such as genic annotations from ENCODE, GENCODE, and Roadmap Epigenomics project provided by this tool^24^. *GARFIELD* evaluates enrichment using generalised linear regression models that account for allele frequency, distance to the nearest gene TSS, and LD. LD correlation based on the UK10K dataset is also provided by the software. In each experimental condition, we used P-values for all SNPs tested in *cis-*eQTL analysis. If a SNP was tested for association with multiple genes, the smallest P-value was kept. Enrichment odds ratios were calculated at various eQTL significance thresholds: 1×10^−3^, 1×10^−4^, …, 1×10^−8^.

### Response eQTL detection

Response eQTLs (reQTLs) were identified in monocytes and T cells separately. For each cell type, we focused on top eSNPs of eGenes that were significant in either resting or stimulated conditions. For eGenes that were significant in both conditions and for which two top eSNPs were not in high LD (r^2^<0.8), we tested both of the top eSNPs; on the other hand, if the two top eSNPs were in high LD, we tested the more significant one, to reduce tests on redundant SNPs. In monocytes, 417 interaction tests involving 398 eGenes were performed, and 1,959 tests involving 1,749 eGenes were performed in T cells. Gene expression data in two conditions were combined within each cell type, and the following linear mixed-effects model was tested for eGene–top eSNP pairs using the *lmer* function in the *lme4* R package^77^:

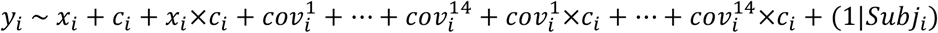

where *y*_*i*_ indicates the expression level of an eGene for the *i*^th^ sample, *x*_*i*_ the SNP allele dosage, *c*_*i*_ the condition (resting: 0 and stimulated: 1) in which the gene expression was measured, 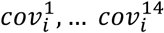 the 14 covariates used in the original eQTL mapping (gender, 3 genotype PCs, and 10 PEER factors), and *subj*_*i*_ the individual from which the *i*^th^ sample was taken. The term *x*_*i*_×*c*_*i*_ models the interaction between the genotype and the condition, and (1|*subj*_*i*_) indicates the individual-specific random effect for this paired study design.

We applied permutations to estimate empirical P-values for the interaction term. In each permutation step, the condition variable was shuffled within each individual, and the same linear mixed model was tested to get the permuted statistics for the interaction term. The permutation adjusted P-value for each interaction test was calculated as follows:

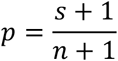

where *n* was the total number of permutations (1,000) and *s* was the number of cases where the permutated statistics were more significant than the original observed ones. We added 1 to both the numerator and the denominator to avoid underestimating permutation P-values^78^. BH FDR-controlling procedure was applied to permutation adjusted P-values and significant interactions were identified at 5% FDR.

### *Trans*-eQTL identification

To detect *trans*-acting genetic regulation of gene expression in each condition, we tested for associations between SNPs and genes that were located on different chromosomes using the same linear model and covariates as in the *cis*-eQTL mapping. We tried the following different approaches to deal with the multiple testing:

#### (1) Genome-wide FDR correction

BH FDR-controlling procedure was applied to nominal P-values from all *trans*-association tests, and significant *trans*-associations were identified at 5% FDR^6^.

#### (2) Gene-level FDR correction

For each gene, P-value of the top SNP was multiplied by 1×10^6^, which was the estimated number of independent SNPs across the genome (calculated as 0.05 divided by the commonly used genome-wide significance threshold of 5×10^−8^). To control gene-level FDR, BH FDR-controlling procedure was then applied to the minimum adjusted P-values for all genes.

#### (3) Gene-level Bonferroni correction

Bonferroni correction was used to control the gene-level FDR, by using a significance P-value threshold of 3.8×10^−12^ (5×10^−8^/13,109, where the denominator indicates the number of genes). The Bonferroni correction was extremely conservative because the tests (or genes) were not independent with each other.

In resting monocytes as well as in LPS-stimulated monocytes, one *trans*-eQTL signal was significant in all three methods. At 5% genome-wide FDR level, we observed 10 and 15 eGenes with significant *trans*-eQTLs (*trans*-eGenes) in resting and PHA-stimulated T cells, respectively, corresponding to a nominal P-value threshold of 1.9×10^−10^ in resting T cells and 2.5×10^−10^ in PHA-treated T cells. The number of significant *trans*-eGenes dropped, respectively, to 6 and 8 by using the gene-level FDR correction (corresponding to a nominal P-value threshold of 5.3×10^−12^ in both conditions), and to 5 and 7 by using the gene-level Bonferroni correction. The limited power was the major issue given the sample size; thus we used genome-wide FDR correction, the least conservative method to determine significant *trans*-eQTLs used in the downstream analysis^6^.

### Mediation analysis

We hypothesised that *trans*-eQTLs regulated the expression of distant genes through *cis-*mediators, or local genes whose expression was regulated by the same *trans*-eQTLs. To test this hypothesis, we focused on the *trans*-eQTLs that were also associated with adjacent *cis-*eGenes, meaning that the *trans*-eQTLs were also *cis-*eQTLs. For each *trans*-eGene–*cis-*eGene pair, we tested the *trans*-eSNP with the smallest P-value as the exposure, a *cis-*eGene as the mediator, and a *trans*-eGene as the outcome (Figure 3B). In total, we tested 14 mediation trios: 1 from resting monocytes, 1 from LPS-stimulated monocytes, 9 from resting T cells, and 3 from PHA-stimulated T cells.

We performed mediation test for the 14 trios using the *mediation* R package^79^. The effect of the exposure on the mediator (C) was estimated in *cis*-eQTL mapping. The effect of the mediator on the outcome (<) adjusting for the exposure and the effect of the exposure on the outcome (2′) adjusting for the mediator were estimated in the following multiple regression:

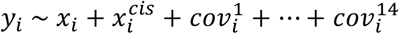

where *y*_*i*_ indicates the value of the outcome (or the expression level of the *trans*-eGene) for the *i*^th^ sample, *x*_*i*_ the exposure (or the eSNP allele dosage), 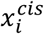 the mediator (or the *cis*-eGene expression), and 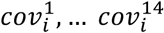 the 14 covariates used in eQTL mapping. The estimates of *b* and *c′* were beta coefficients for 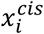 and *x*_*i*_, respectively. The “direct effect” of the exposure on the outcome was quantified as *c′*, the “indirect effect” of the exposure on the outcome through the mediator was quantified as C×<, and the “total effect” was the sum of the previous two effects^80^. Complete mediation occurs when the direct effect *c′* is zero after controlling for the mediator, and partial mediation happens when the direct effect is different from zero. To identify significant mediation trios (the null hypothesis *H*_0_:*ab* = 0), we used a nonparametric bootstrap method (10,000 simulations) implemented in the *mediation* R package for variance estimation and P-value calculation. BH FDR-controlling procedure was applied to correct for multiple testing.

### Genetic overlap of eQTLs and disease

We downloaded publicly-available GWAS data from the following three resources: ImmunoBase (**URLs**), LD Hub^81^ (**URLs**), and GWAS Catalog^82^ (**URLs**). If one disease was investigated in multiple studies, we focused on the most recent one, which usually had the largest sample size. We also downloaded summary statistics of GWAS that were carried out using the ImmunoChip array from the ImmunoBase resource; this array was designed for immunogenetics studies and captured more variants of immune-relevant genetic loci^83^. We had summary statistics for the following immune-mediated diseases: allergic disease (asthma, hay fever, or eczema)^84^, allergic rhinitis^85^, allergic sensitisation^85^, asthma^86^, childhood-onset asthma^41^, adult-onset asthma^41^, inflammatory bowel disease (IBD) including its two subtypes – Crohn’s disease and ulcerative colitis^87^, celiac disease^88,89^, autoimmune thyroid disease^90^, juvenile idiopathic arthritis^91^, multiple sclerosis^92^, narcolepsy^93^, primary biliary cirrhosis (PBC)^94,95^, primary sclerosing cholangitis (PSC)^96^, psoriasis^97^, rheumatoid arthritis^98,99^, systemic lupus erythematosus (SLE)^100^, and type 1 diabetes^101,102^. These datasets contained summary statistics obtained using European populations for both significant and non-significant genetic variants, and GRCh37 genomic coordinates were available.

We performed enrichment tests for each of the four sets of significant eQTLs using *GARFIELD*^24^. Generalised linear models were applied to test for enrichment in eQTLs of variants associated with the above diseases at a significance threshold of 1×10^−6^. Bonferroni correction was applied to correct for multiple testing, where the number of tests was the number of GWAS datasets (24) multiplied by the number of eQTL datasets (4), and the Bonferroni-adjusted P-value threshold was 5.2×10^−4^, adjusting for the 4×24 (96) tests.

### Colocalisation of *cis-*eQTLs with disease associations

We applied a Bayesian method implemented in the *coloc* v3.1 R package^26^ to test whether any of the disease-associated GWAS loci shared the same causal variants with the *cis*-eQTLs. Full summary statistics were required to run the colocalisation analysis using *coloc*. For loci where *cis-*eQTLs were also associated with diseases at a P-value threshold of 1×10^−6^, colocalisation test was performed on a 400-kb window centered on the top *cis*-eSNP. For each locus, colocalisation test was performed on overlapping SNPs where both eQTL and GWAS summary statistics were available. We excluded regions where not enough SNPs (<25) were available for colocalisation test. As shown in Guo *et al*.^103^, selection of different prior probabilities of a SNP being causal for both of the traits had effects on the posterior support for colocalisation. To be conservative, we used a lower prior probability of 1×10^−6^ instead of the default value of 1×10^−5^.

For each locus, the Bayesian method assessed the support for the following five exclusive hypotheses: no causal variants for either of the two traits (H_0_), a causal variant for gene expression only (H_1_), a causal variant for disease risk only (H_2_), distinct causal variants for two traits (H_3_), and the same shared causal variant for both traits (H_4_). The package estimated posterior probabilities (PP_0_, PP_1_, PP_2_, PP_3_, PP_4_) to summarise the evidence for the above five hypotheses. High PP_1_ or PP_2_ and low PP_3_ + PP_4_ indicate a lack of power to identify the causal signals^26^. We excluded loci where PP_3_ + PP_4_ <0.8, and focused on loci with strong evidence support for shared causal variants (H_4_), i.e. ratio of PP_4_ to PP_3_ ≥5.

### Mendelian randomisation analysis

To investigate causal effects of eGene expression on the above immune-mediated diseases, we performed a two-sample Mendelian randomisation (MR) analysis. Summary statistics from both our eQTL and external GWAS studies were required, including beta coefficient and its standard error, effective allele (based on which the beta was estimated), the other allele, and P-value. For each disease trait, we used *cis-*eSNPs that were also included in the GWAS dataset, and removed ambiguous variants (if any) using the *TwoSampleMR* R package^104^. We then selected LD pruned (r^2^ <0.1) *cis-*eSNPs as genetic instrumental variables (IVs). We focused on eGenes with at least 3 genetic IVs available, and performed the following MR methods implemented in the *MendelianRandomization* R package^105^: inverse variance weighted (IVW), weighted median, weighted mode, and MR Egger. These methods have different assumptions for valid IVs: IVW assumes that all IVs are valid^106^; weighted median assumes that valid IVs contribute to more than 50% of the weight^107^; weighted mode assumes that the largest group of IVs are valid^108^; MR Egger regression, which is the least sensitive, assumes that the pleiotropic effects of IVs are not correlated with the genetic effects on exposure^109^. We excluded the causal associations for which the intercept in the MR Egger method was significantly not equal to 0, indicating significant average pleiotropic effects. Gene expression was considered to have suggestive evidence of causal effects when at least 3 out of the 4 methods provided significant P-value (≤0.05). In the **Results** section, we reported the statistics of the weighted mode method, which has the least assumption among all methods except the MR Egger, but more sensitive than MR Egger.

## URLs

DICE, https://www.dice-database.org

ImmunoBase, https://www.immunobase.org/

LD Hub, http://ldsc.broadinstitute.org/ldhub/

GWAS Catalog, https://www.ebi.ac.uk/gwas/

