## Supplemental Figures for "Neonatal genetics of gene expression reveal the origins of autoimmune and allergic disease risk"

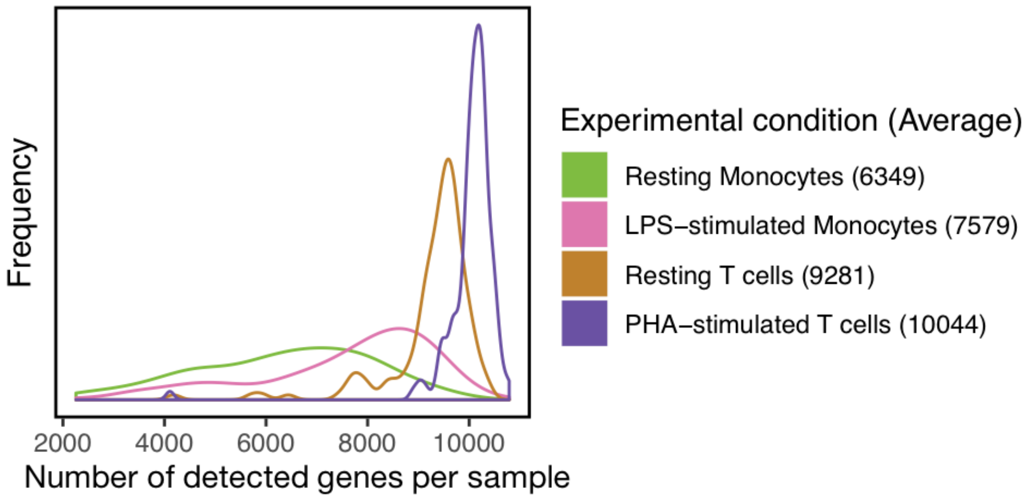


**Figure S1: Distribution of the number of detected genes per microarray sample.** Colour indicates four experimental conditions. Number in brackets indicates the average number of detected genes (**Methods**) within each group.

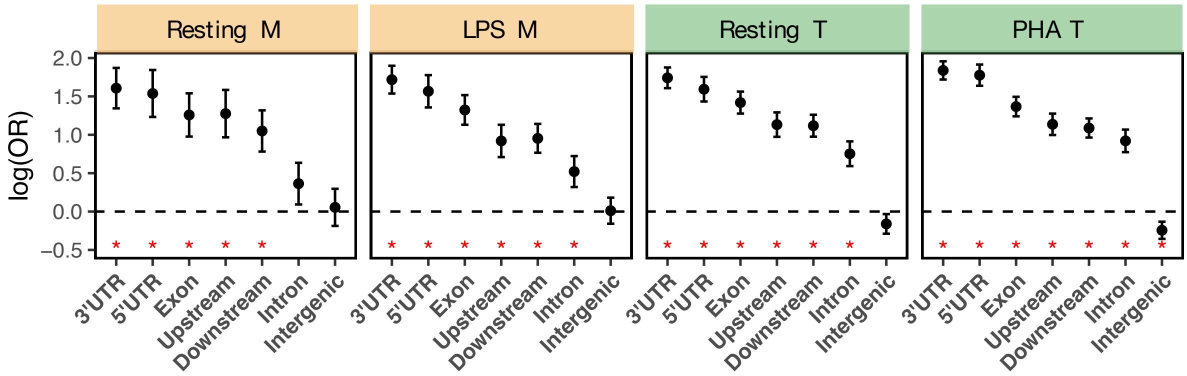


**Figure S2: Enrichment of *cis*-eQTLs in 3'UTR, 5'UTR, and exon regions.** Four plots show the enrichment of *cis*-eQTL SNPs identified in four experimental conditions ("M" and "T" indicate monocytes and T cells, respectively). Logarithm of Odds Ratio (OR), estimated at the eQTL significance level of $1\times10^{-5}$, is shown on y-axes (**Methods**), and the 95% confidence intervals are plotted. Tests that were significant after the Bonferroni correction are marked by red asterisks, where a P-value threshold of 0.0018 was used, adjusting for the 7×4 (28) tests.

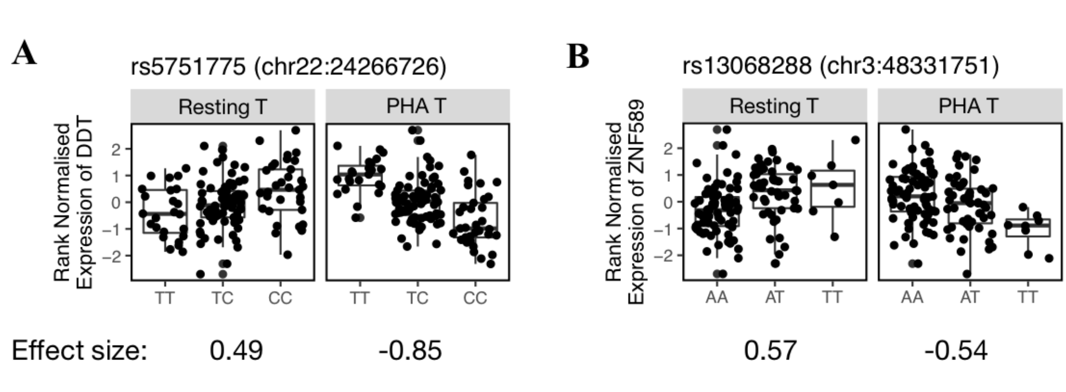


**Figure S3: Response eQTLs (reQTLs) with flipped directions of effects on gene expression across conditions.** ReQTLs of *DDT* (**A**) and *ZNF585* (**B**) show opposite directions of eQTL effects. (**A**) Two boxplots show associations of rs5751775 (top eSNP in resting T cells) with *DDT* expression in resting (left) and PHA-stimulated (right) T cells. Dots indicate individuals stratified by genotypes (x-axes) and the rank-normalised gene expression is shown on y-axes. Similarly, panel (**B**) shows the associations of rs13068288 (top eSNP in PHA-stimulated T cells) with *ZNF589* expression. Chromosome number and genomic position in hg19/GRCh37 are in brackets next to the SNP rsID. Effect sizes of these associations are shown under each box plot, which is the number of s.d. of gene expression increased per allele dosage.

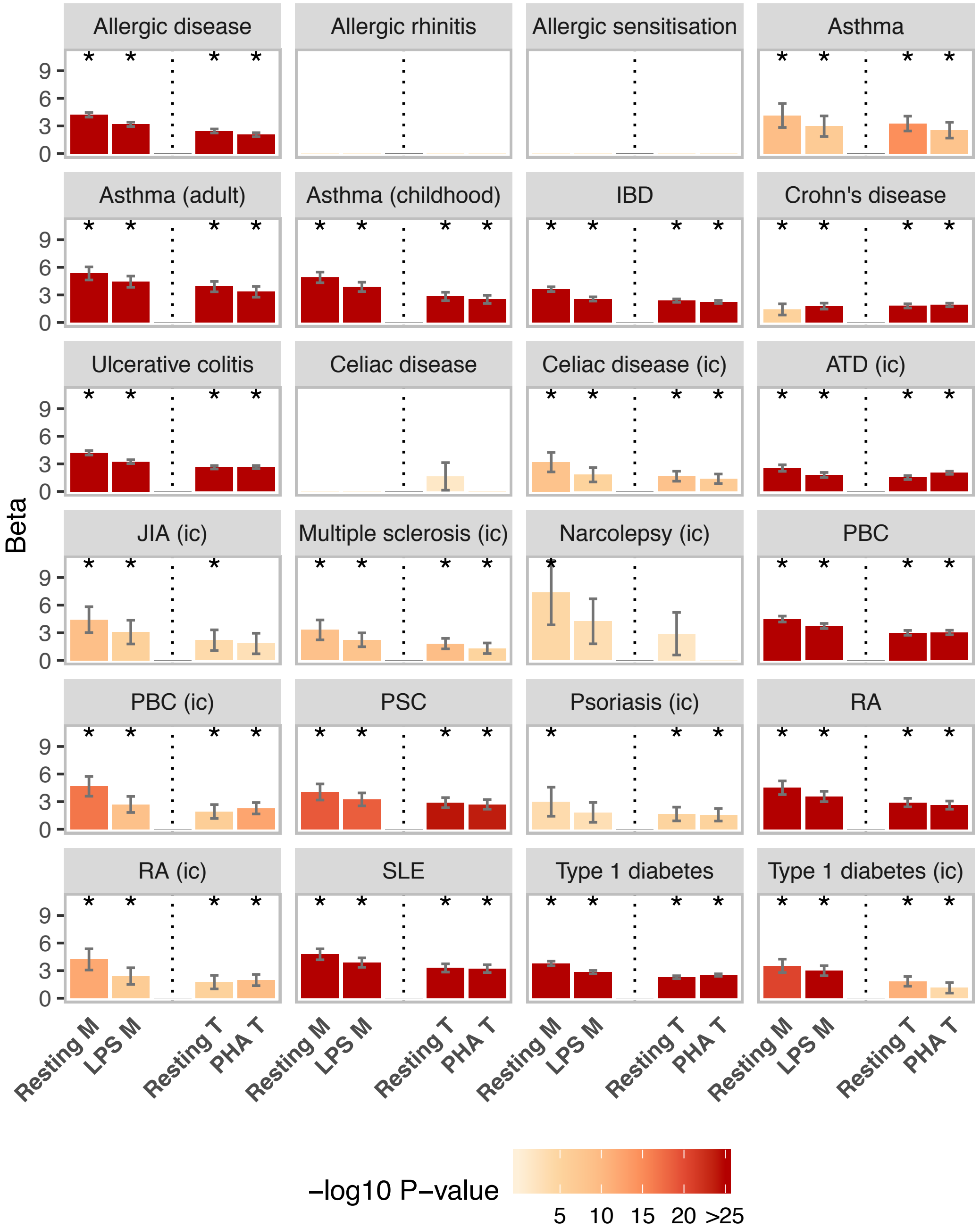


**Figure S4: Enrichment of early-life *cis*-eQTLs for genetic variants associated with immune-related diseases.** Each plot represents one disease, with "ic" indicating that the study was performed using ImmunoChip array (**Methods**). Disease abbreviations are as follows: "IBD": inflammatory bowel disease, "ATD": autoimmune thyroid disease, "JIA": juvenile idiopathic arthritis, "PBC": primary biliary cirrhosis, "PSC": primary sclerosing cholangitis, "RA": rheumatoid arthritis, and "SLE": systemic lupus erythematosus. Enrichment was tested for each of the four sets of *cis*-eQTLs separately (x-axes: "M" and "T" indicate monocytes and T cells, respectively). Height of each bar indicates the beta coefficient (or log of the odds ratio) from enrichment analysis, with 95% confidence intervals (CIs) shown as the error bars. Tests that are not significant at P-value 0.05 are not shown since the beta estimates are not reliable and the CIs are too large. Colour of each bar indicates the significance level based on minus log10 P-values; the same colour is used if the P-value is smaller than $1\times10^{-25}$. Asterisks indicate that the tests are significant after Bonferroni correction for multiple testing.

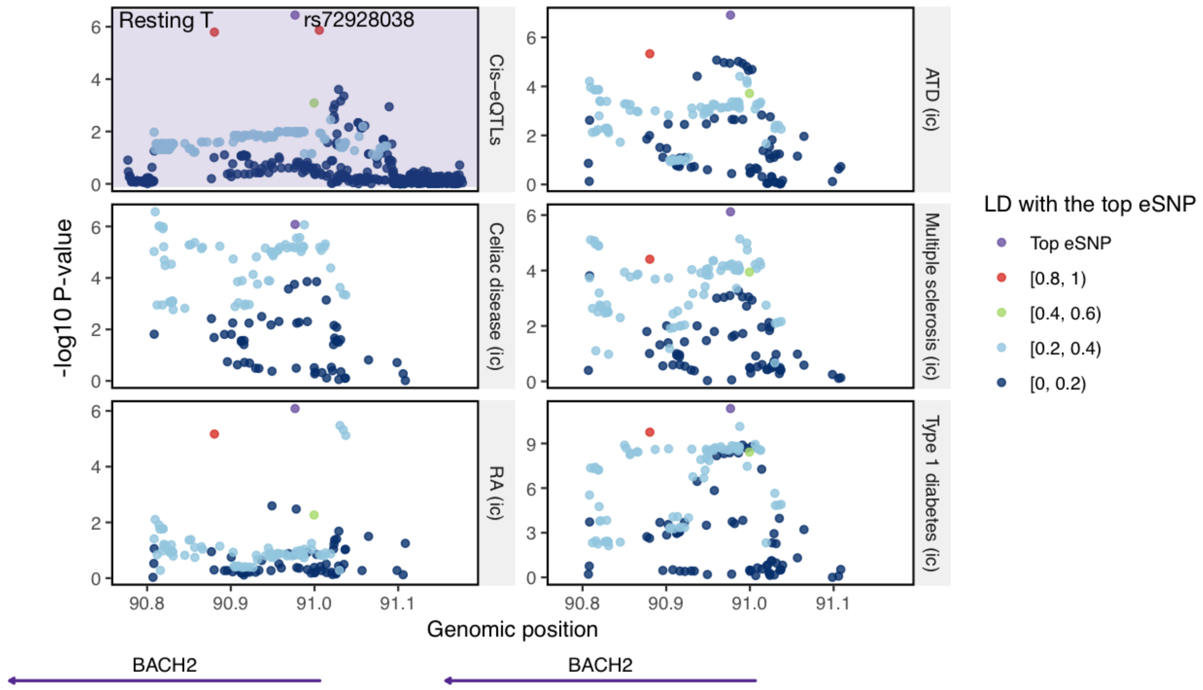


**Figure S5: Colocalisation of the *cis*-eQTL of *BACH2* with multiple diseases.** Six regional plots show eQTL association with *BACH2* expression in resting T cells (purple background), and GWAS associations with autoimmune thyroid disease (ATD), celiac disease, multiple sclerosis, rheumatoid arthritis (RA), and type 1 diabetes (white background). "ic" indicates that the GWAS was performed using ImmunoChip array. The minus log10 P-value is plotted on y-axes for all SNPs located within 200 kb from the top eSNP of *BACH2* (rs72928038). Colours of dots indicate the LD correlation with the top eSNP (purple). Positions of genes located on this locus are shown at the bottom. *BACH2* did not have significant *cis*-eQTLs in the other three experimental conditions.

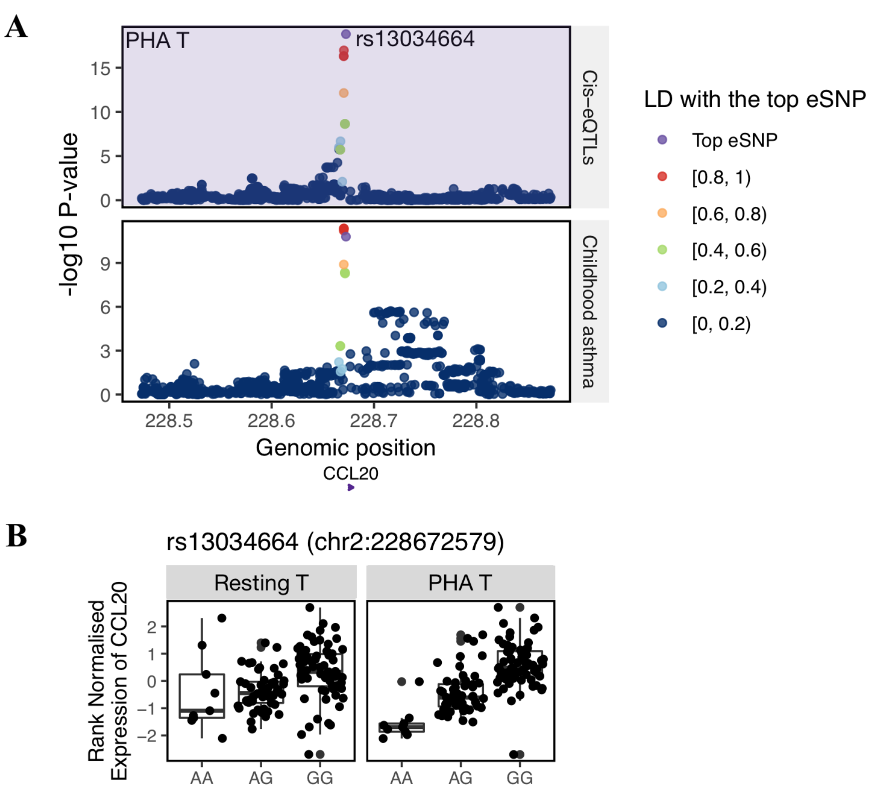


**Figure S6: Colocalisation between the response eQTL (reQTL) of *CCL20* and childhood-onset asthma association.** (**A**) Regional plots show eQTL association with *CCL20* expression in PHA-stimulated T cells (purple background), and GWAS association with childhood-onset asthma (white background). The minus log10 P-value is plotted on y-axes for all SNPs located within 200 kb from the top eSNP of *CCL20* (rs13034664). Colours of dots indicate the LD correlation with the top eSNP (purple) based on CAS genotype data. Positions of genes located on this locus are shown at the bottom. *CCL20* did not have significant *cis*-eQTLs in the other three experimental conditions. (**B**) Box plots show the rank-normalised *CCL20* expression (y-axes) in resting (left) and stimulated (right) T cells stratified by genotypes of the reQTL (x-axes). In resting T cells, no SNP was significantly associated with *CCL20*.

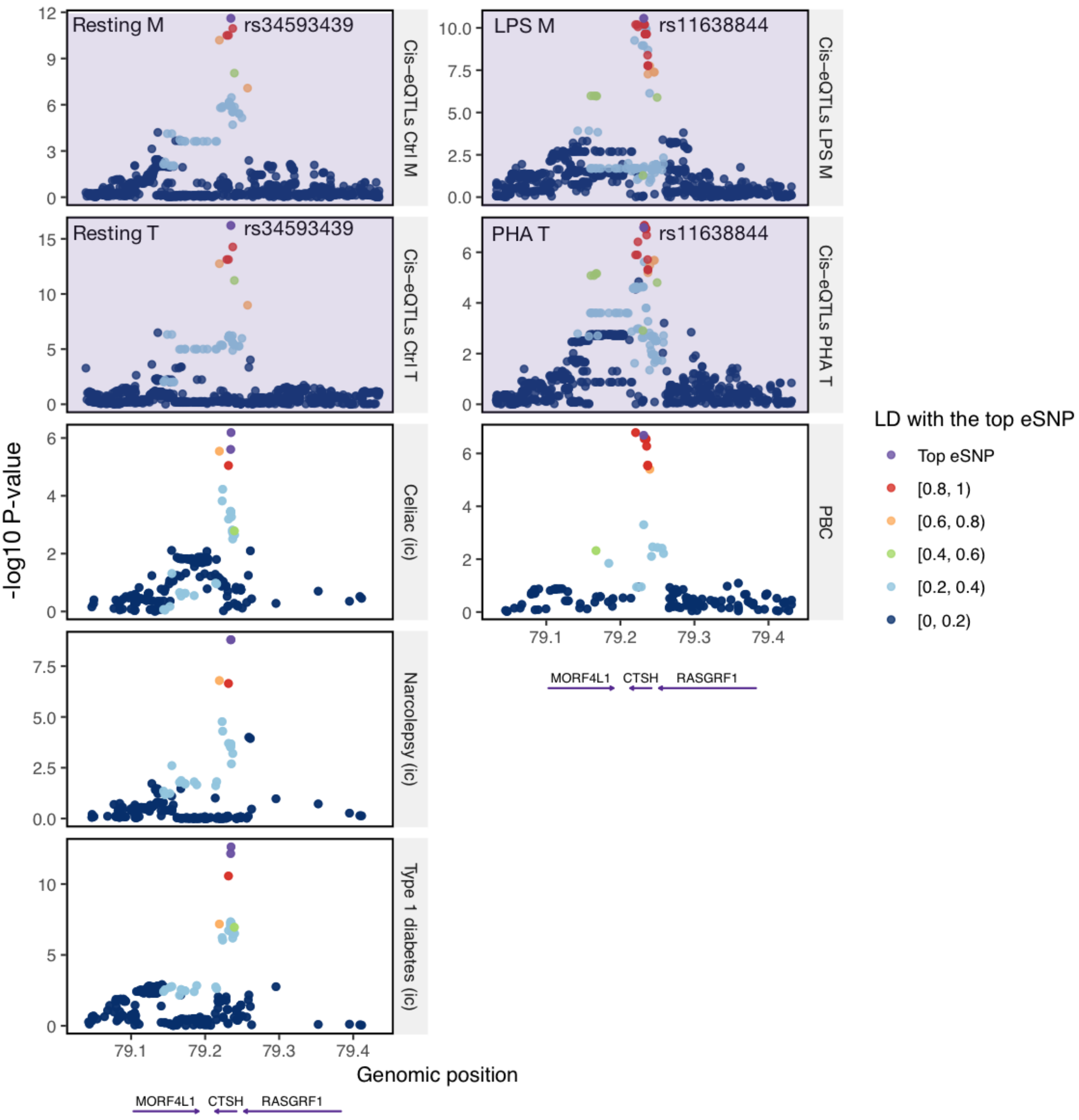


**Figure S7: Colocalisations with different diseases are observed for the *CTSH cis*-eQTLs identified in resting (left) and stimulated (right) cells**. On the left, regional plots show eQTL associations (purple background) with *CTSH* expression identified in resting monocytes (Resting M) and resting T cells (Resting T). The two eQTL signals have the same top eSNP rs34593439, and both colocalised with GWAS hits associated with celiac disease, narcolepsy, and type 1 diabetes (white background). "ic" indicates that the GWAS was performed using ImmunoChip array. On the right, regional plots show eQTL associations (purple background) with *CTSH* expression identified in LPS-stimulated monocytes (LPS M; top eSNP rs11638844) and PHA-stimulated T cells (PHA T). They both colocalised with GWAS signal for primary biliary cirrhosis (PBC; white background). The minus log10 P-value is plotted on y-axes for all SNPs located within 200 kb from the corresponding top eSNP (left: rs34593439; right: rs11638844). Colours of dots indicate the LD correlation with the top eSNP (purple). Positions of genes located on this locus are shown at the bottom.

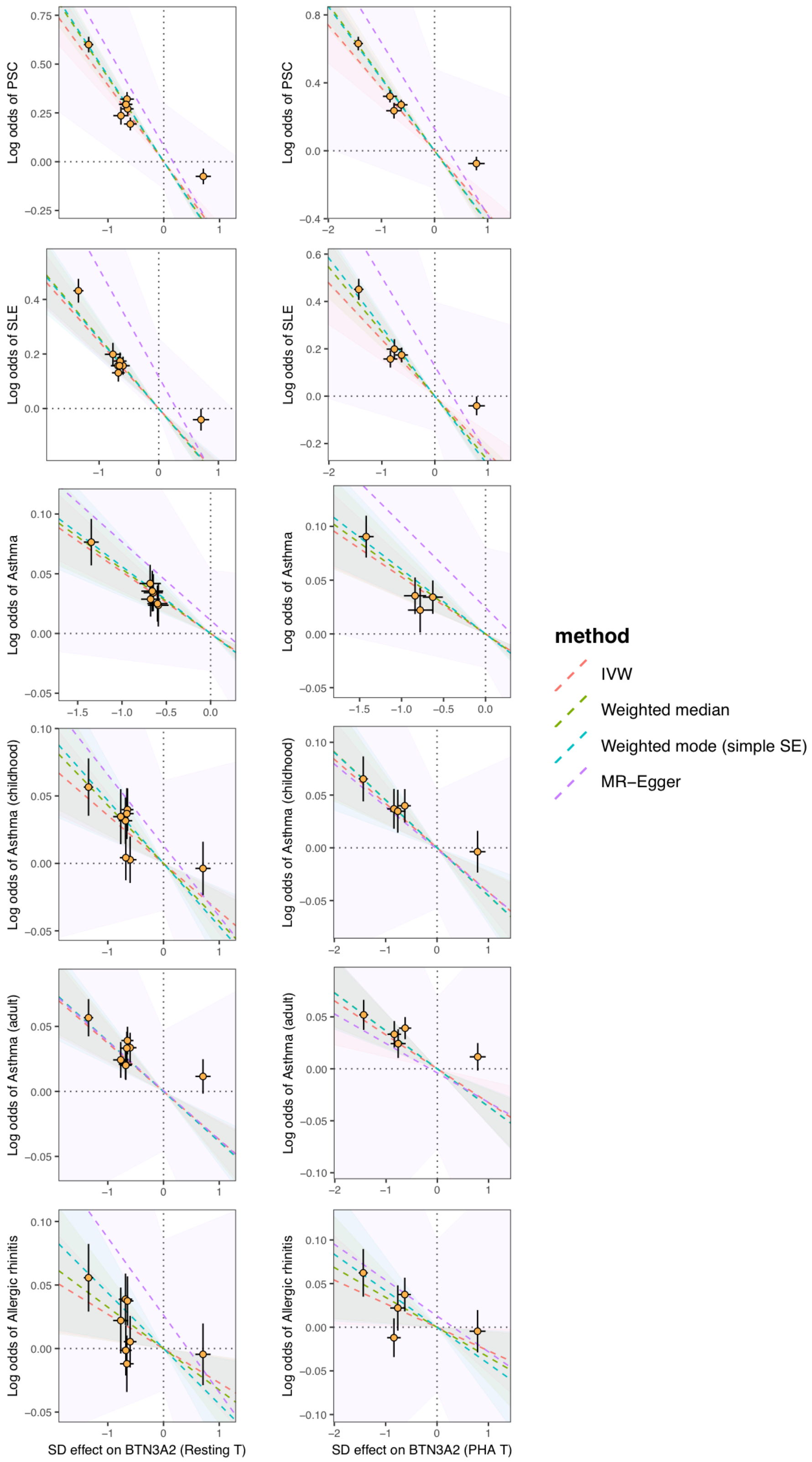


**Figure S8: Causal associations (negative) between *BTN3A2* expression and immune-related diseases.** *Cis*-eQTL effect sizes with 95% confidence interval (CI; x-axes) estimated from resting T cells (left) and PHA-stimulated T cells (right) are plotted against the genetic effect sizes with 95% CI on disease risk (y-axes). Rows show immune-related diseases that were negatively associated with *BTN3A2* expression (PSC: primary sclerosing cholangitis; SLE: systemic lupus erythematosus). In each plot, four fitted lines indicate different Mendelian randomisation methods and ribbons indicate their 95% confidence intervals (IVW: inverse variance weighted; **Methods**).

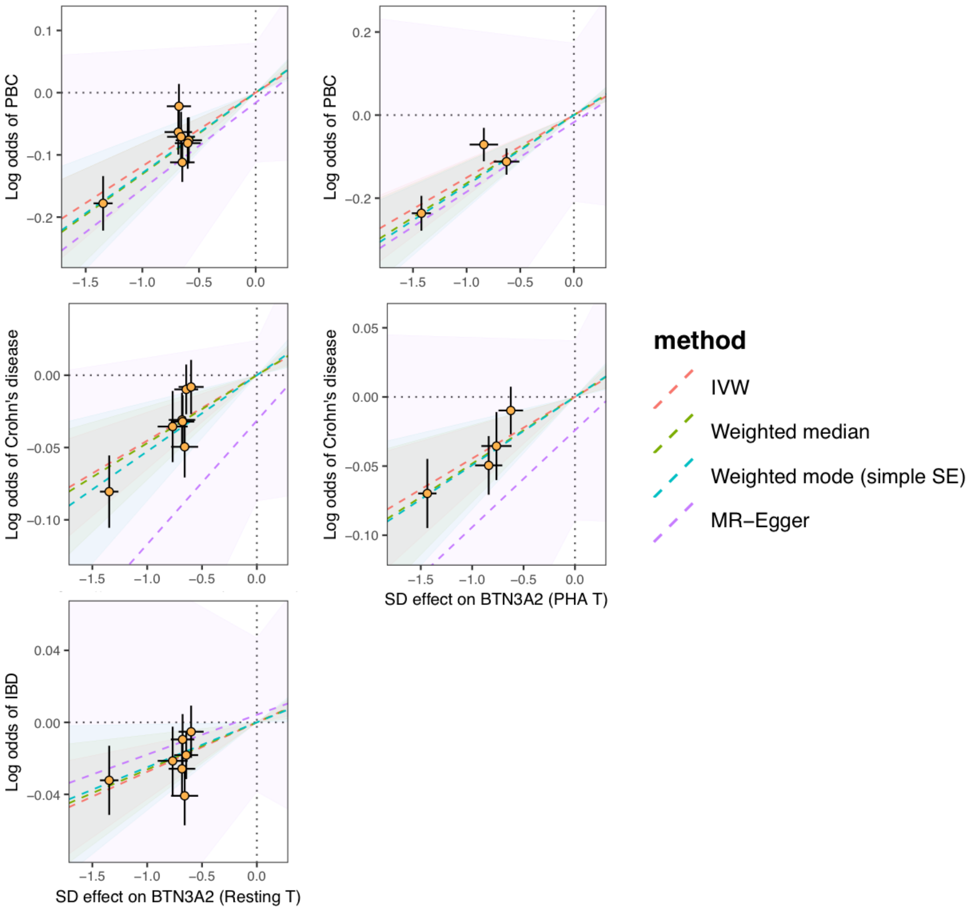


**Figure S9: Causal associations (positive) between *BTN3A2* expression and immune-related diseases.** *Cis*-eQTL effect sizes with 95% confidence interval (CI; x-axes) estimated from resting T cells (left) and PHA-stimulated T cells (right) are plotted against the genetic effect sizes with 95% CI on disease risk (y-axes). Rows show immune-related diseases that were positively associated with *BTN3A2* expression (PBC: primary biliary cirrhosis; IBD: inflammatory bowel disease). In each plot, four fitted lines indicate different Mendelian randomisation methods and ribbons indicate their 95% confidence intervals (IVW: inverse variance weighted; **Methods**).

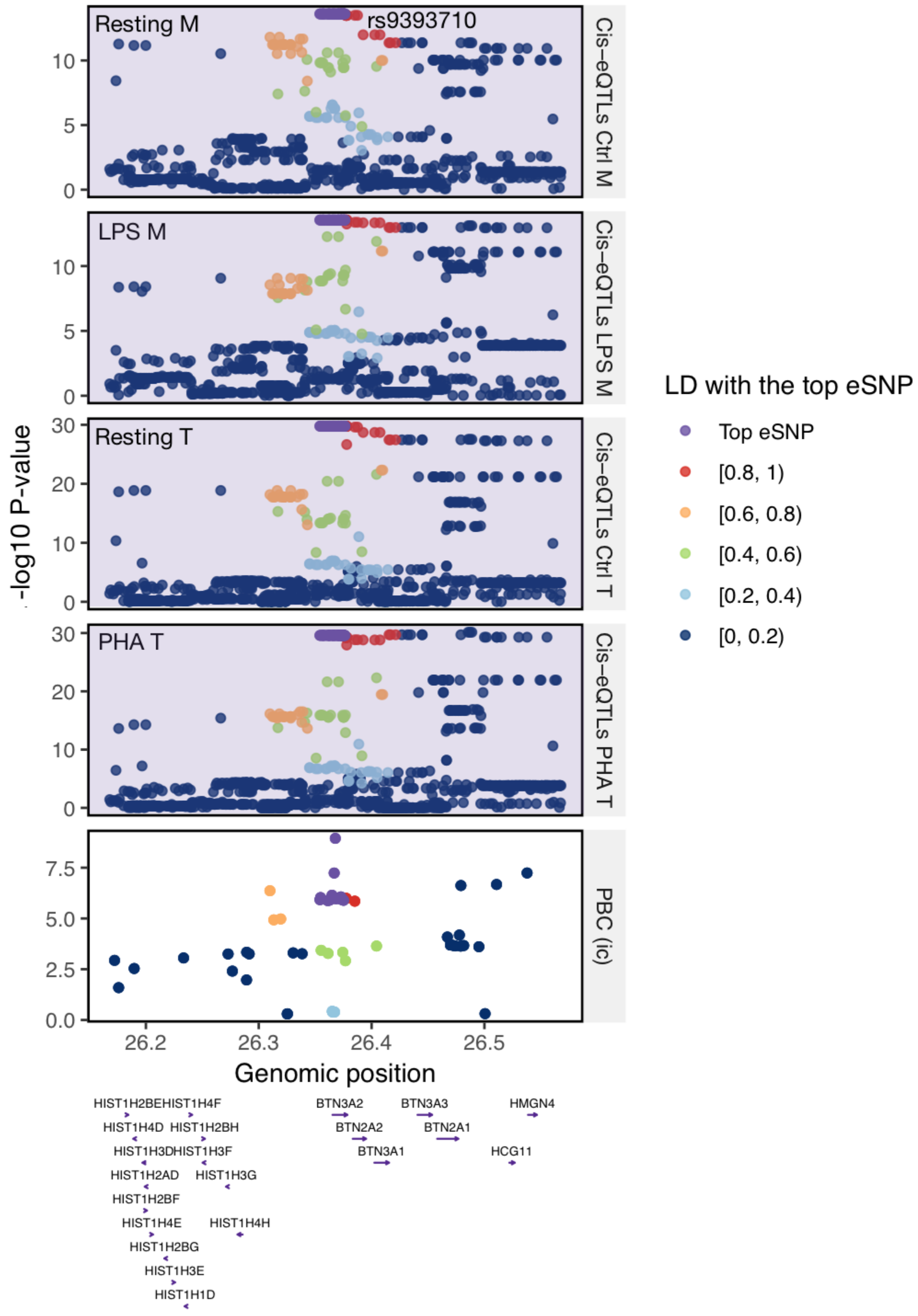


**Figure S10: Colocalisation between the *cis*-eQTLs of *BTN3A2* in four experimental conditions with primary biliary cirrhosis (PBC).** Five regional plots show eQTL associations (purple background) with *BTN3A2* expression in four conditions ("M" and "T" indicate monocytes and T cells, respectively), and GWAS association with PBC (white background). "ic" indicates that the GWAS was performed using ImmunoChip array. The minus log10 P-value is plotted on y-axes for all SNPs located within 200 kb from rs9393710, which was the top SNP of the GWAS hit. rs9393710 was among the top eSNPs identified in resting and stimulated monocytes and in resting T cells, and the top eSNP in stimulated T cells was in high LD with it ($r^2 = 0.92$). Colours of dots indicate the LD correlation with the top eSNP (purple) based on CAS genotype data. Positions of genes located on this locus are shown at the bottom.

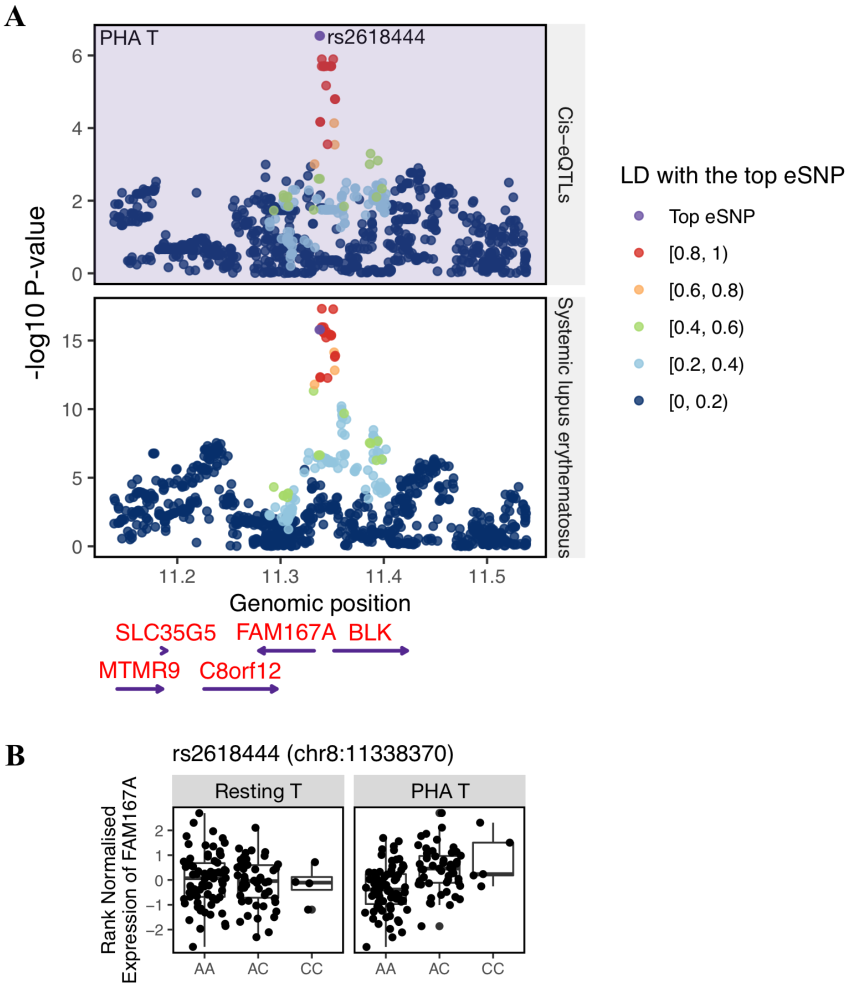


**Figure S11: Colocalisation between the response eQTL (reQTL) of *FAM167A* and systemic lupus erythematosus.** (**A**) Regional plots show eQTL association with gene expression of *FAM167A* in PHA-stimulated T cells (purple background), and GWAS association with systemic lupus erythematosus (white background). The minus log10 P-value is plotted on y-axes for all SNPs located within 200 kb from the top eSNP of *FAM167A* (rs2618444). Colours of dots indicate the LD correlation with the top eSNP (purple). Positions of genes located on this locus are shown at the bottom. (**B**) Box plots show the rank-normalised *FAM167A* expression (y-axes) in resting T cells (left) and in PHA-stimulated T cells (right) stratified by genotypes of the reQTL rs2618444 (x-axes). In resting T cells, no SNP was significantly associated with *FAM167A*.
